# Cone photoreceptor ablation in microglia-deficient larval zebrafish retina elicits a regenerative response alongside a compensatory immune cell response

**DOI:** 10.64898/2026.02.26.708140

**Authors:** Jordan E Rumford, Ashley A Farre, Justin Mai, Halle V Weimar, Claire D Shelton, Michael Morales, Diana M Mitchell

**Affiliations:** Department of Biological Sciences, University of Idaho, Moscow, ID, USA

**Author notes:** Author for Correspondence: Diana M. Mitchell, Associate Professor, Biological Sciences 875 Perimeter Drive, MS 3051 University of Idaho, Moscow, ID U.S.A. 83844-3051.

**Keywords:** microglia, retina, regeneration, zebrafish, cone photoreceptors, inflammation, Müller glia

## Abstract

Emerging evidence implicates retinal microglia and inflammation as important components impacting the outcome of retinal regeneration, which is spontaneously achieved in zebrafish retina following acute damage but is limited or blocked in mammals. In this paper, we describe the regenerative response in the larval zebrafish retina following ablation of cone photoreceptors. To investigate the role of microglia in the regenerative response, we used both *irf8^st95^* heterozygote (microglia-sufficient) and *irf8^st95^* homozygous mutant (microglia-deficient) zebrafish. We compared multiple aspects of the regenerative response in *irf8*+/− and *irf8*−/− larval retinas, including entry of the Müller glia (MG) into the cell cycle, the amplification of MG-derived progenitor cell (MGPC) proliferation, inflammatory and glial reactivity-associated gene expression, and the regeneration of cones. We found only modest impacts to early and late stages of MGPC proliferation and to inflammatory gene expression in *irf8* mutants, with no obvious impacts to the regeneration of cones. Notably, we detected a population of immune cells in *irf8* mutants that emerged following cone ablation, which expanded in number then were reduced over time, following a trajectory similar to microglia-sufficient siblings but at markedly reduced abundance. The immune cells detected in *irf8* mutants included a subset with L-plastin/4C4 antibody staining patterns different than those in microglia-sufficient siblings, suggesting distinct origins and/or phenotype compared to resident microglia in controls. Though strong conclusions about the role of microglia were limited due to the presence of such immune cell populations in *irf8* mutants, our results are consistent with several reports that indicate a role for microglia and inflammation in regulating MGPC proliferation in the regenerating retina. Collectively considered with other reports, our results further indicate that compensatory responses, which may include different immune cells and/or signaling from other retinal cell types such as the Müller glia, are elicited in microglia-deficient retinas upon neuronal damage.

## Introduction

The zebrafish retina has a remarkable and well-appreciated intrinsic regenerative capacity. Because the cell types involved are also present in mammalian retinas, work in zebrafish has generated important translatable knowledge about the cells and genetic pathways that result in successful regeneration of retinal neurons. In zebrafish retina, Müller glia (MG) respond to acute damage of neurons by dividing asymmetrically [1] to produce a neuronal progenitor cell pool that then amplifies in abundance via proliferation [2–6]. The MG-derived progenitor cells (MGPCs) subsequently differentiate into neurons that integrate into retinal circuits to restore visual function [7–12]. Temporally alongside (or even prior to) the Müller glial response, microglia also respond to acute retinal damage [13–21]. These features of glial responses have been demonstrated in numerous experimental systems of neuronal damage to the zebrafish retina [1, 3, 6, 7, 14, 18, 21–24]. Several studies indicate that inflammatory signals, to which microglia likely contribute, are important for Müller glial responses and MGPC proliferation [14, 15, 21, 25, 26] and may also impact later outcomes such as the survival of regenerated neurons [21, 27, 28]. Given their activation in multiple retinal degenerative diseases, acute retinal injury [13, 17, 27, 28], and intercellular interactions with the Müller glia [29, 30], there is strong interest in identifying the function of microglia in retinal regeneration as this will lead to a better understanding of supportive versus detrimental activities of these cells which could be modulated therapeutically. Microglia could provide inflammatory signals and cues that play into a broad signaling axis [31, 32]. Alternatively, or in addition, their role as phagocytes for clearance of debris and damaged/dying cells [13, 29, 33] could shape regenerative responses that follow neuronal death.

The first published studies to probe the role of microglia in retinal regeneration in zebrafish used distinct tools to modulate microglial presence in the retina upon acute damage. These included a transgenic system in which the *mpeg1* promoter (active in zebrafish microglia and macrophages [34, 35]) drives bacterial nitroreductase (NTR) conferring sensitivity to the pro-drug metronidazole [14], a pharmacological approach for CSF1R inhibition using the drug PLX3397 [20, 36], and a genetic mutant with deficient microglial development [36]. Further, the immunosuppressive drug dexamethasone has been used to probe immune signaling in retinal regenerative responses [14, 15, 21, 22, 26, 36] and though such experiments suggest inflammatory signaling may be involved they do not directly implicate microglia because this drug has broad effects on multiple cell types including the Müller glia [37, 38]. A more recent paper used nanoparticle technology to target dexamethasone to microglia, which increased kinetics of rod photoreceptor regeneration [39]. The generation of a zebrafish line with a loss-of-function mutation in the gene *irf8* provided a novel genetic model for microglia deficiency [40] and is an attractive tool to further probe the role of microglia in retinal regeneration. To date this zebrafish *irf8* mutant has been utilized in several studies focusing on the retina [15, 18, 36, 41]. Other zebrafish genetic mutants have since been generated which are deficient in microglia; these include *csf1r* mutants though the two zebrafish paralogs encoding *csf1ra* and *csf1rb* have differential impact on microglial development [42]. To sufficiently deplete microglia, *csf1ra/b* double mutants may be required, which may require extensive genetic crosses to generate the necessary fish for experiments.

Currently, the literature does not provide a clear consensus about the role of microglia in retinal regeneration. While some studies in zebrafish suggest retinal regeneration is hindered in the absence of microglial functions [15, 20, 31, 36, 43] other work was less supportive of such a role [18]. Some of the underlying factors that may have resulted in different conclusions between these reports could potentially be the differences in microglia abundance or phenotype in the different mutant lines (*irf8* vs *csf1ra/b*), use of different drug-based approaches, differences in the retinal damage systems used, or the age of the fish used in the experiments (larval versus adult). Timing of microglial inhibition may also be a factor [14, 39]. Further, in mouse retina with forced neuronal production after damage, microglia were found to inhibit retinal regeneration [28], suggesting microglia could behave differently in different species.

Given our previous work demonstrating effects on Müller glia (MG) in microglia-deficient *irf8^st95^* mutant zebrafish [41], an interest to further probe the role of microglia in retinal regeneration, and the ability to achieve microglial deficiency with a single gene mutation, we examined MG-mediated regenerative responses and the regeneration of cone photoreceptors in the larval zebrafish retina in the *irf8^st95^* mutant. We used metronidazole (Mtz)-mediated ablation of cone photoreceptors in transgenic *gnat2*:nfsb-mCherry [23, 44] fish at larval ages. Our results are consistent with several components of multiple reports investigating the role of microglia in retinal regeneration in zebrafish [14, 15, 18, 36]. Homozygous *irf8* mutants showed only modest decreases in MGPC proliferation and inflammatory gene expression following cone ablation, while regeneration of cones at 7- and 14-days post injury was not significantly impacted in *irf8* mutants. While *irf8* mutants were highly deficient in microglia in larval retina at baseline, upon induced retinal damage (i.e. ablation of cones), leukocytes consistent with microglia/macrophage features were detected in *irf8* mutants causing difficulties in conclusive connections about the role of microglia in regeneration of cones. Expression patterns of selected microglia markers commonly used in zebrafish (4C4 and L-plastin antibody labeling), as well as leukocyte localization in the damaged and regenerating retina, were different in *irf8* mutants compared to heterozygous siblings. Collectively, our results suggest that compensatory innate immune responses, which may include immune cell populations phenotypically distinct from resident microglia and/or signaling from other retinal cell types such as the Müller glia, are elicited in microglia-deficient retinas upon photoreceptor damage.

## Results

### Ablation of cones in larval zebrafish retina activates microglial responses

Using *gnat2*:nfsb-mCherry transgenic zebrafish larvae [23], in which the Nfsb/NTR enzyme tagged with mCherry fluorescent protein is expressed in cone photoreceptors, we administered DMSO (vehicle) or the pro-drug metronidazole (Mtz) at ∼4.5 days post fertilization (dpf) via immersion to induce death of cones. After 24-48 hours of treatment, larvae were collected for analysis of cone death and glial responses (Figure 1). Although ablation of a central patch of cones has been reported in *gnat2*:nfsb-mCherry larvae with shorter Mtz treatment duration [23], we aimed to ablate all cones in our experiments and found that this required longer duration of Mtz immersion. We did not detect reliable, consistent ablation of cones in larval retinas with 24-hour Mtz exposure. However, by 48 hours of Mtz treatment, mCherry+ cones showed morphological alterations and disorganization indicating collapse compared to DMSO treatment (Fig 1A, B). Retinal cryosections further recapitulated this result at 48 hours post-treatment with Mtz (48 hpt) and additionally demonstrated abundant TUNEL staining in the outer nuclear layer (ONL) that co-localized with collapsed mCherry (cone reporter) signal, indicating that cones were ablated with this treatment regime (Fig 1C,D). Microglia were visualized by staining with the 4C4 antibody, thought to recognize LGALS3BP [45] and commonly used to detect zebrafish microglia. 4C4+ microglia were rarely detected within the ONL of DMSO treated retinas (Fig 1C) but were readily seen within the damaged ONL of Mtz treated retinas at 48 hpt (Fig 1D). The microglia were positioned around and within dying cone signal and often observed to encircle TUNEL+ signal (Fig 1D, D’, D’’), consistent with microglial engulfment of dying cones. In addition, some mCherry and TUNEL signal was occasionally detected in the INL (Fig 1D), which is consistent with phagocytic capacity of the Müller glia [29, 41] and previous reports of MG engulfing photoreceptor debris upon damage [46].

**Figure 1:**
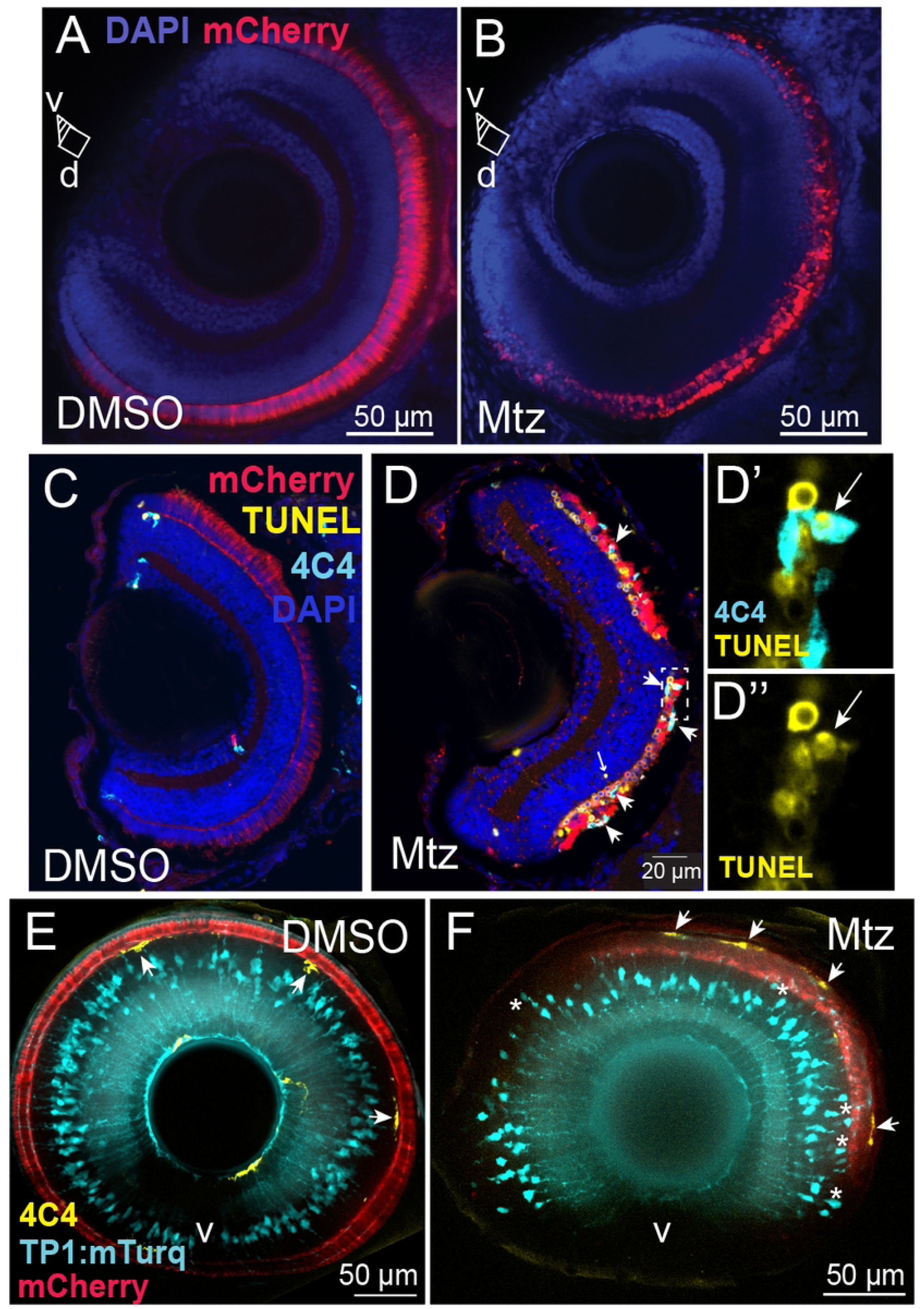
Ablation of cones in *gnat2*:nfsb-mCherry larvae and early glial responses. A, B. Whole larval eyes imaged on intact *gnat2*:nfsb-mCherrry zebrafish larvae, using confocal microscopy, at 48 hours post treatment (hpt) with DMSO (A) or metronidazole (Mtz, B) visualizing nuclear (DAPI) stain and mCherry fluorescence in cones. Annotation v, d indicate ventral and dorsal, respectively, with ventral located at the bottom of the image stack and dorsal at top. C, D. Retinal cryosections collected at 48 hpt following DMSO (C) or Mtz (D) treatment, and stained for microglia (4C4 antibody), TUNEL, and DAPI; mCherry fluorescence from cones. Microglia localized to collapsed, TUNEL+ cones in Mtz treated retinas (arrowheads, D) and occasional TUNEL/mCherry signal is seen in the inner retina (arrow, D). D’, D’’ is an enlarged region (dotted box in D) showing microglia (4C4, cyan) engulfing TUNEL+ cones (yellow) in Mtz treated retinas. E, F. Whole eyes from larval *gnat2*:nfsb-mCherry;TP1:mTurquoise double transgenic fish, stained for microglia (4C4), and imaged by confocal microscopy at 24 hpt (DMSO or Mtz treatment). Selected optical sections are projected showing retina radially around the lens. Arrows indicated microglia localized at basal face of ONL/OPL in DMSO treated samples (E) and their transition to the apical region of the ONL in Mtz treated samples (F). Asterisks (*) in F show Müller glia cell bodies that have shifted apically towards the ONL.

Examining whole eyes/retinas 24 hours earlier (at 24 hpt), we observed that microglia showed differential localization by 24 hpt with Mtz compared to DMSO (Figure 1E,F). Microglia were identified at the OPL/interface of the ONL in DMSO samples (Figure 1E) but in Mtz treated retinas, microglia were localized to the apical regions of the ONL (Figure 1F). We used the *TP1*:mTurquoise reporter to visualize Müller glia (MG) [29], which revealed that in addition to changes in microglial localization by 24 hours of Mtz treatment, MG also showed changes. Positioning of MG cell bodies upon Mtz treatment was altered in comparison to DMSO controls with several MG bodies localizing more apically towards the ONL (Fig 1E,F). Collectively, these images indicate that 48 hours of Mtz treatment reliably induces the death of cones in *gnat2*:nfsb-mCherry larval fish. Further, changes observed for microglia and Müller glia indicate that these cell types sense changes to cones by 24 hpt.

We next examined microglia over time following cone ablation, by staining retinal cryosections from *gnat2*:nfsb-mCherry fish collected at 24-hour intervals from 48-144 hours post-treatment (Fig 2). Again, at 48 hpt we observed responding microglia interacting with mCherry+ cone signal in the damaged ONL upon Mtz treatment (Fig 2B). We observed microglia localized to the ONL at later timepoints (Fig 2C,D), and microglia were observed adjacent to mCherry+ cone reporter signal in retinal sections from larvae at 96-144 hpt Mtz (Fig 2C,D). Quantification of microglia numbers over time, using 4C4 antibody to label microglia, revealed that microglial numbers were increased by 48 hpt following Mtz treatment and remained elevated compared to DMSO samples through at least 144 hpt (Fig 2G). Collectively, the localization patterns and microglial numbers in retinal cryosections suggest that microglia could perform functions involved with MG responses and regeneration of cones.

**Figure 2:**
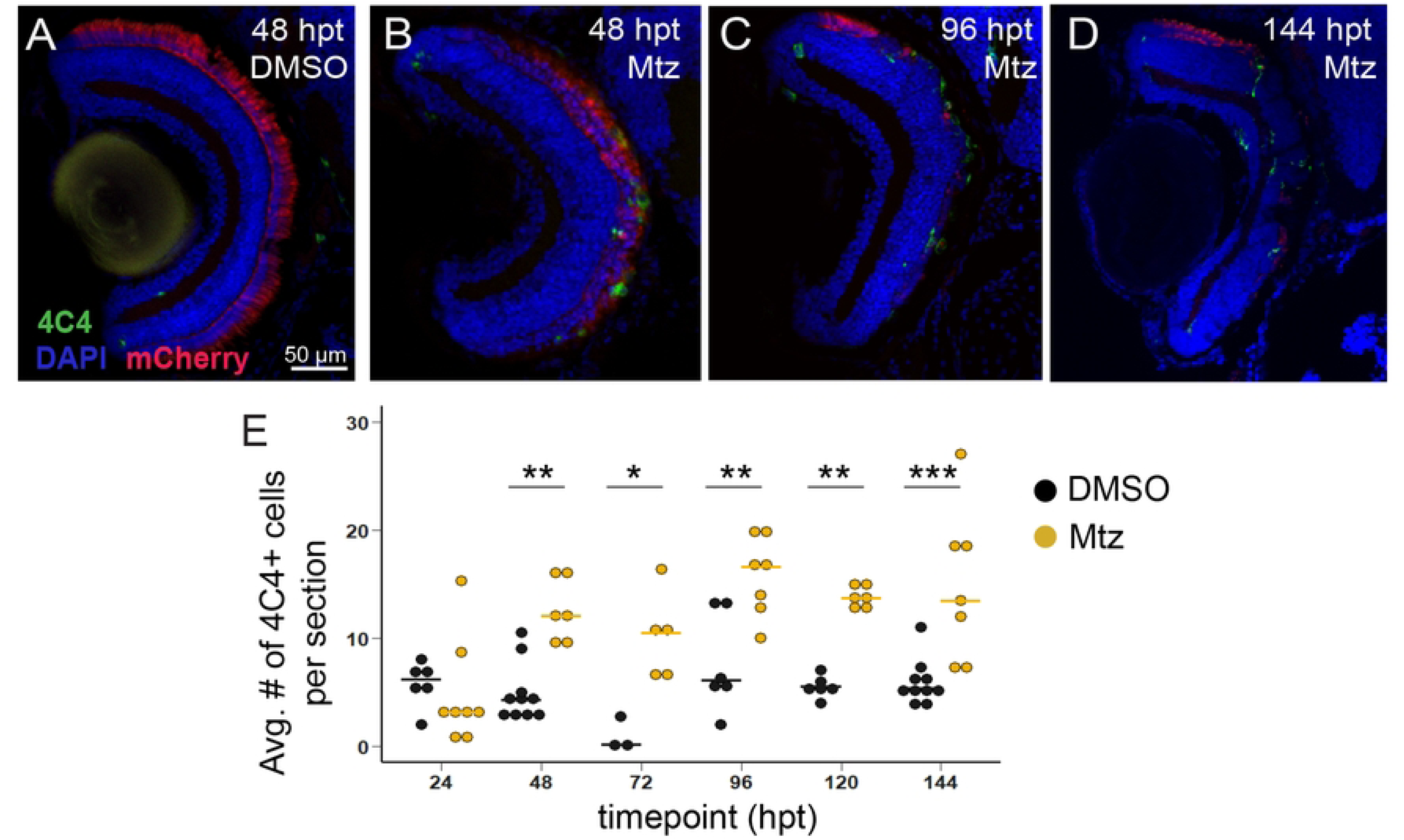
Microglia response over time following cone ablation in *gnat2*:nfsb-mCherry larval retinas. A-D. Retinal cryosections from *gnat2*:nfsb-mCherry fish collected at the indicated timepoints (hours post treatment, hpt) following DMSO (A) or Mtz treatment (B-D) and stained for 4C4 (microglia) and DAPI. Cones visualized by mCherry fluorescence. Scale bar in A applies to all images. E. Quantification of numbers of 4C4+ microglia per retinal cryosection at the indicated timepoints following DMSO or Mtz treatment. Two-way ANOVA indicated an effect of treatment and timepoint (p<0.05); Tukey’s post-hoc comparisons are shown between DMSO and Mtz groups for each timepoint, *p<0.05, **p<0.01, ***p<0.001.

### Entry of Müller glia into the first asymmetric division is not altered in *irf8* mutants

Previous work has shown that following retinal damage in zebrafish, the responding Müller glia enter an initial asymmetric division to produce a pool of proliferating progenitor cells [1]. To begin to dissect the role of microglia in regeneration of cones in our system, we first examined the timing of the initial Müller glial asymmetric division following Mtz treatment of *gnat2*:nfsb-mCherry larvae. We collected samples at 24 and 48 hpt and stained retinal sections for glutamine synthetase (GS) and PCNA to label Müller glia that had entered the cell cycle (Fig 3A-C). At 24 hours following DMSO or Mtz treatment, GS+PCNA+ cells were not significantly detected in the inner nuclear layer (INL) (Fig 3B, 3D). At 48 hpt, numerous GS+PCNA+ (Müller glia in S-phase) were detected in the INL of Mtz treated samples compared to DMSO treated samples (Fig 3C, 3D), indicating that Müller glia enter the cell cycle between 24 and 48 hours following Mtz treatment. We further confirmed the timing of MG cell cycle entry using EdU immersion. In these experiments, DMSO/Mtz treatment followed the same schedule, with EdU added from 24-48 hpt. Retinal sections from samples collected at 48 hpt were stained for GS, PCNA, and EdU (Fig 3E-G). Quantification of EdU incorporation revealed that by 48 hpt, a subset of PCNA+ Müller glia also incorporated EdU to detectable levels (Fig 3H) while there were essentially no Müller glia that stained positive for EdU but negative for PCNA (Fig 3I, which would indicate MG that had completed a cell cycle). These results indicate that the first responding Müller glia enter S-phase within the window of 24-48 hpt following Mtz treatment and that this window labels Müller glia in the first asymmetric division.

**Figure 3:**
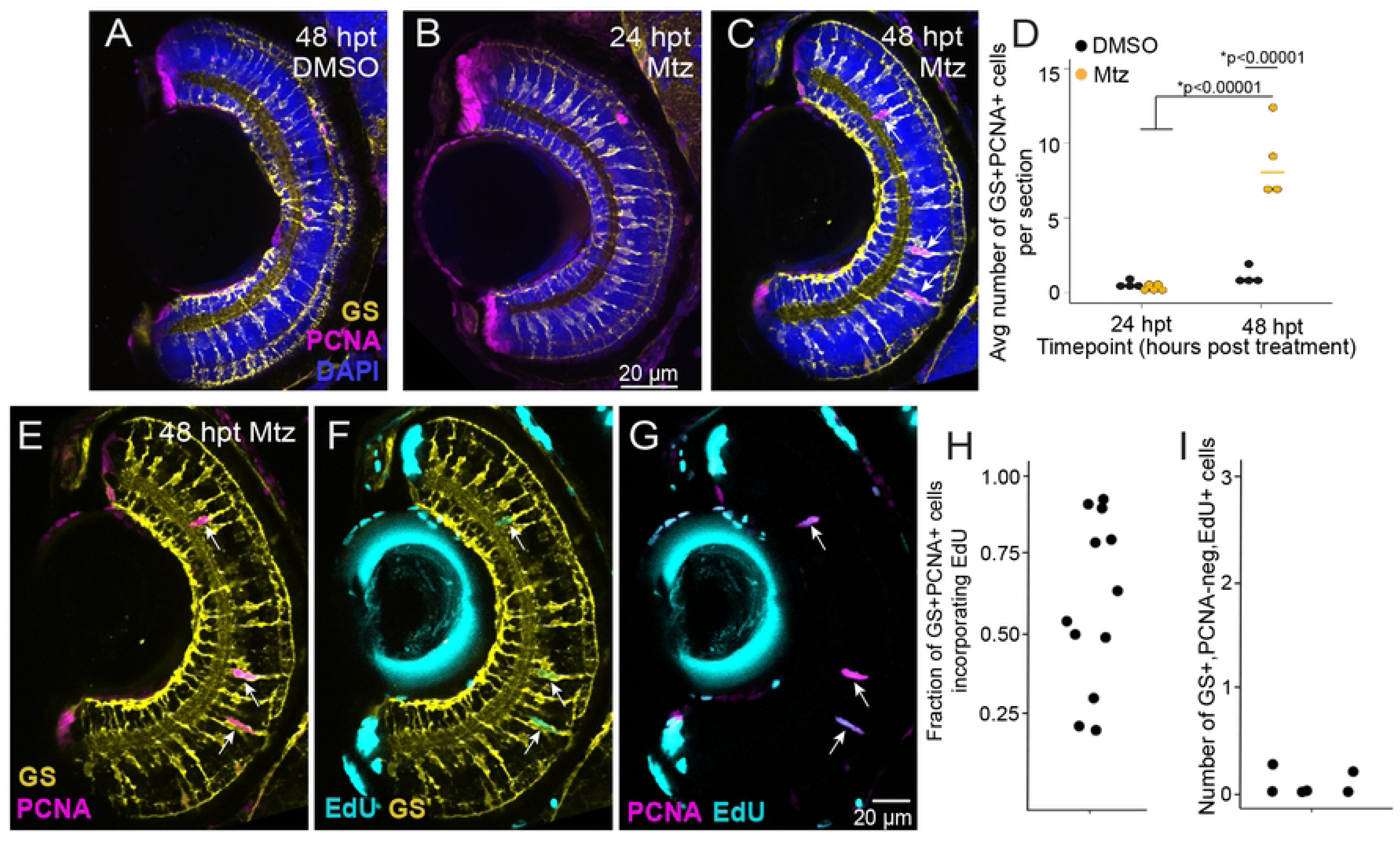
Müller glia enter the first asymmetric division between 24 and 48 hpt following Mtz treatment of *gnat2*:nfsb-mCherry larvae. Representative images of retinal cryosections from *gnat2*:nfsb-mCherry fish collected at the indicated timepoints (hours post treatment, hpt) following DMSO (A) or Mtz treatment (B,C), and stained for Glutamine Synthetase (GS), PCNA (Proliferating Cell Nuclear Antigen) and DAPI. Scale bare in B applies to A-C. C. Arrows indicate GS+PCNA+ Müller glia in the inner nuclear layer (INL). D. Quantification of the number of GS+PCNA+ cells per retinal section following DMSO or Mtz treatment at the indicated timepoints. **p<1×10^-5^, 2-way ANOVA with Tukey’s post-hoc. E-G. Representative images of retinal cryosections from *gnat2*:nfsb-mCherry fish collected at 48 hpt of Mtz treatment, following immersion in EdU from 24 to 48 hpt. Images show staining and detection of GS, PCNA, and EdU; arrows indicate GS+ Muller glia in the INL that are also PCNA+ and EdU+. Scale bar in G applies to E, F, G. H. The fraction of GS+PCNA+ cells that were also EdU+ detected in retinal cryosections collected at 48 hpt following Mtz treatment and exposed to EdU from 24 to 48 hpt. I. The number of GS+ cells that were PCNA-negative but EdU+ in the cryosections collected at 48 hpt.

To determine if the first asymmetric MG division upon cone ablation is altered in *irf8* mutant larvae compared to siblings, we crossed *gnat2*:nfsb-mCherry;*irf8*+/− breeders to *irf8*−/−fish to produce offspring carrying the *gnat2*:nfsb-mCherry transgene (for cone ablation) that are either microglia-sufficient (*irf8*+/−) or microglia-deficient (*irf8*−/−). We and others have shown microglia are deficient in numbers in *irf8*−/− larval retinas at timepoints prior to and spanning the Mtz treatment window in our experiments [36, 41]. At 48 hpt following DMSO or Mtz immersion, larvae were collected for retinal cryosections. We performed microglia/leukocyte staining in a subset of samples at 48 hpt (∼6.5 dpf age). This analysis was done because microglia/leukocyte numbers may change over development and/or in response to DMSO/Mtz treatment. Further, peripheral macrophages are reported to develop in *irf8* mutants around 6 days of age [40] (which coincides with our 48 hpt timepoint) and could potentially respond to cone damage. We examined retinal sections after staining with both 4C4 antibody and an antibody to L-plastin (Lcp1), a pan-leukocyte marker in zebrafish [13, 47] (Fig 4). In *irf8*+/− retinas, cells that stained for both Lcp1 and 4C4 were readily identified in the damaged cone layer (Fig 4A-A’’). In *irf8*−/−retinas, the number of cells staining for these markers was significantly reduced though a limited number of Lcp1+ cells (sometimes but not always co-labeling with 4C4) were detected in the damaged ONL (Fig 4B-B’’). Quantifications of Lcp1+ and 4C4+ cell numbers revealed significantly reduced numbers in *irf8*−/− compared to *irf8*+/− siblings (Fig 4C,D); however, it is worth noting that some cells staining with these markers were indeed detected in *irf8* mutant retinal sections. Therefore, the *irf8* mutant remains deficient in microglia/leukocyte numbers in response to cone damage at 48 hpt. We noted that a lower percentage of the Lcp1+ cells detected in *irf8* mutants also co-labeled with the 4C4 microglia marker compared to *irf8*+/− (Fig 4E), suggesting that the leukocytes detected in *irf8* mutants may be phenotypically different than the responding microglia/macrophages in *irf8*+/−.

**Figure 4:**
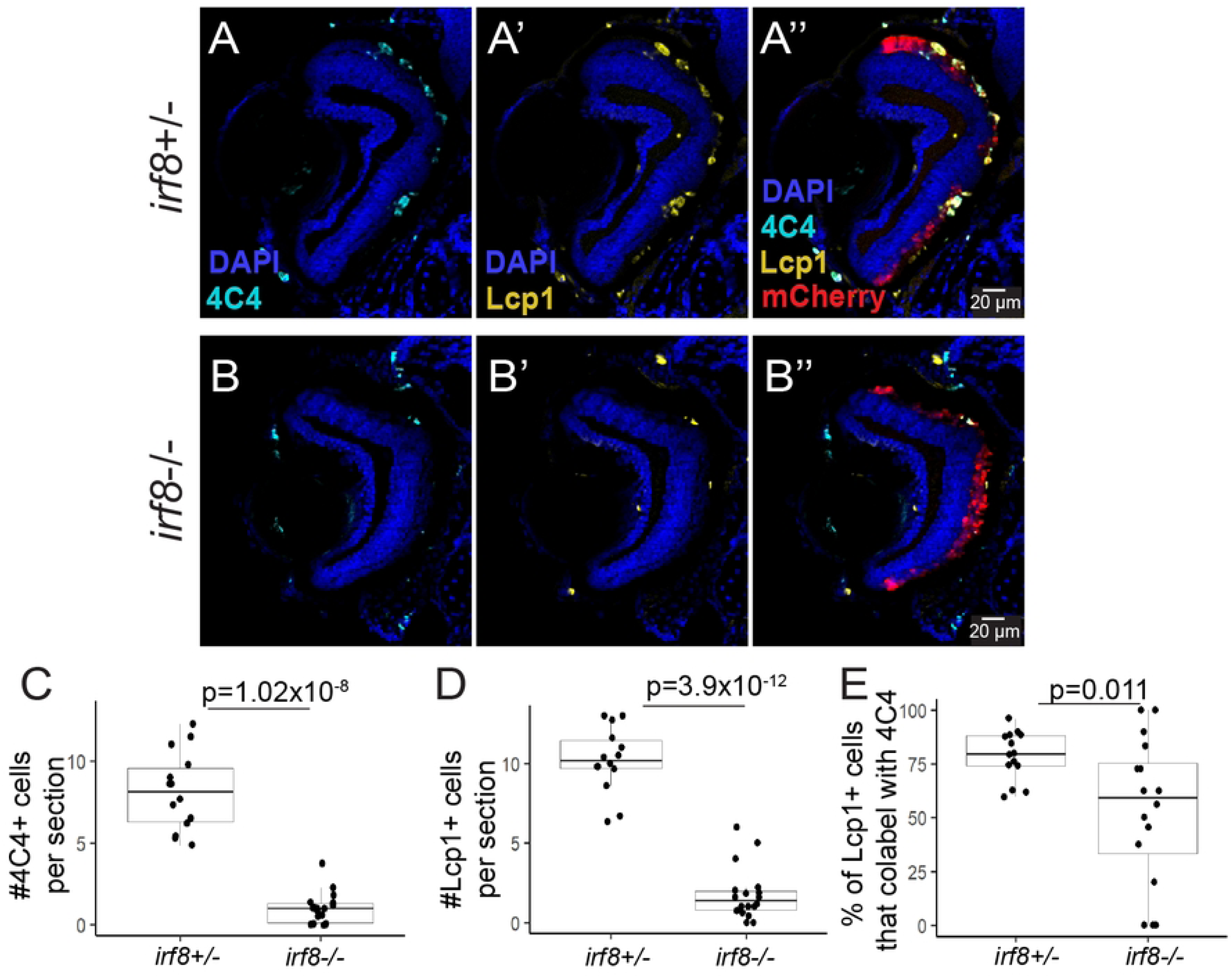
Microglia are reduced in number in *gnat2*:nfsb-mCherry;*irf8* mutant compared to *irf8* heterozygote retinas at 48 hpt following cone ablation. Retinal cryosections from *gnat2*:nfsb-mCherry;*irf8*+/− or *gnat2*:nfsb-mCherry;*irf8*−/− larval zebrafish samples collected at 48 hours post Mtz treatment and stained for 4C4, Lcp1, and DAPI; cones are visualized by mCherry fluorescence. A-A’’. Microglia/leukocytes that stain for Lcp1 and 4C4 are detected in the ONL in response to cone ablation at 48 hpt in *irf8*+/− retinas. Scale bare in A’’ applies to A-A’’. B-B’’. Some, but reduced, Lcp1 and 4C4+ cells are detected in the ONL in response to cone ablation at 48 hpt in *irf8*−/− retinas. Scale bare in B’’ applies to B-B’’. C. Quantification of the number of 4C4+ cells in retinal cryosections of *irf8*+/− or *irf8*−/− fish at 48 hpt following Mtz treatment. D. Quantification of the number of Lcp1+ cells in retinal cryosections of *irf8*+/− or *irf8*−/− fish at 48 hpt following Mtz treatment. E. The percent of Lcp1+ cells that also co-label with 4C4 in the two genotypes. P-values are indicated for pairwise comparisons between *irf8*+/− and *irf8*−/− (Welch’s test).

Given that the *irf8*−/− fish still represent a system of microglia deficiency at 48hpt, we used staining for GS, PCNA, and EdU to analyze the first MG asymmetric division in microglia-sufficient (*irf8*+/) and microglia-deficient (*irf8*−/−) retinal sections from *gnat2*:nfsb-mCherry larvae (Fig 5). Larvae were immersed in Mtz for 48 hours to ablate cones, with EdU addition from 24-48 hpt (Fig 5A). GS+PCNA+ MG were detected in the INL of both genotypes, with some PCNA+ MG also showing EdU incorporation (Fig 5B-C’’’). Comparing the two genotypes, we found no statistically significant differences in the number of MG that stained positive for PCNA at 48 hpt (Fig 5D) nor in the fraction of these PCNA+ MG that had also incorporated EdU at 48 hpt (Fig 5E), indicating that entry of MG into the cell cycle in response to cone ablation occurred similarly by this timepoint in both microglia-sufficient and microglia-deficient conditions. We also quantified the number of MG derived progenitor cells (MGPCs) at 48 hpt by quantifying the number of GS-negative/PCNA+EdU+ cells in the INL (excluding the ciliary marginal zone) (Fig 5F). At this timepoint, MGPCs were not yet abundantly detected in either genotype (Fig 5F) indicating that a pool of abundant proliferating progenitors had not yet accumulated at 48 hpt in both microglia-sufficient and -deficient retinas.

**Figure 5:**
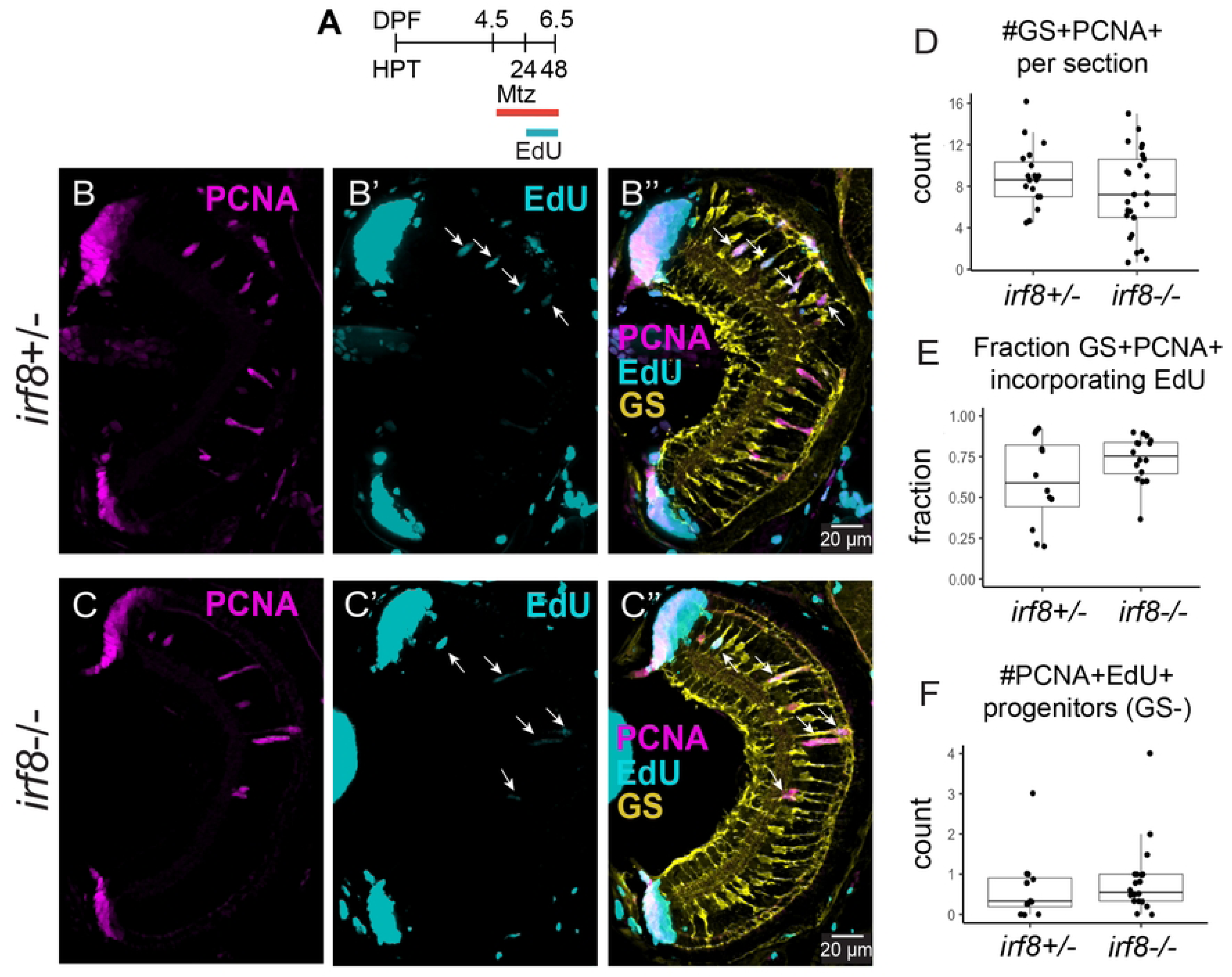
Entry of Müller glia into first asymmetric division following cone death is not significantly impacted in *irf8* mutant retinas. A. Timeline showing treatment of *gnat2*:nfsb-mCherry;*irf8*+/− or *gnat2*:nfsb-mCherry;*irf8*−/− fish with Mtz and exposure to EdU. B-B’’. Images of retinal cryosections from *irf8* heterozygotes collected at 48 hpt, showing PCNA, EdU, and GS signal. Scale bar in B’’ applies to B-B’’. C-C’’. Images of retinal cryosections from *irf8* mutants collected at 48 hpt, showing PCNA, EdU, and GS signal. Arrows in B-C’’ indicate GS+PCNA+EdU+ cells. Scale bar in C’’ applies to C-C’’. D. Quantification of the number of GS+PCNA+ cells per retinal section in the two genotypes. E. The fraction of GS+PCNA+ cells that were also EdU+ in the two genotypes. F. The number of GS-negative, PCNA+EdU+ cells (considered cycling MG-derived progenitors) in the two genotypes. Differences were not statistically significant, Welch’s test.

Given the detection of responding leukocytes in *irf8* mutants (Fig 4), we attempted to inhibit microglial responses with the drug PLX3397. PLX3397 is a CSF1R inhibitor first used to deplete microglia in mouse [48, 49] and more recently has been used for this purpose in zebrafish [20, 36]. Treatment of larvae with PLX3397 (or vehicle) began at 2 dpf and solutions were refreshed every 24 hours. DMSO or Mtz treatment for 48 hours began at 4.5 dpf and EdU was added from 24-48 hpt (Supp Fig1A). In retinal sections from *gnat2*:nfsb-mCherry larvae with no cone ablation (DMSO samples), we detected a reduction in both 4C4+ and Lcp1+ cell numbers with PLX3397 (Supp Fig1B,C) indicating that PLX3397 (PLX) treatment at least partially depletes microglial numbers in undamaged retinas. However, upon cone ablation with Mtz treatment, both 4C4+ and Lcp1+ cell counts were significantly increased resulting in similar overall numbers in both non-PLX and PLX-treated samples (Supp Fig1B,C) indicating that PLX treatment, even continued through 48 hpt/6.5 dpf alongside cone ablation, did not maintain microglial deficiency and/or prevent leukocyte responses to cone damage. Analysis of the first MG cell division did not detect differences in PCNA staining or EdU incorporation by Müller glia in non-PLX and PLX-treated groups at 48 hpt Mtz treatment (Supp Fig1D-G). In these experiments, a small number of proliferating progenitors were detected at 48 hpt (Supp Fig1H) but these did not return statistically significant differences in PLX-treated samples compared to controls. Given the microglial and leukocyte responses induced upon cone damage in PLX treated samples, we could not reliably derive conclusions about microglial functions in these experiments. In addition, this system did not provide a better alternative in terms of microglia/macrophage abundance to that of the *irf8* mutants for probing microglial function in cone regeneration. Therefore, we elected to use only the *irf8* genetic system for further experiments in this paper.

### Inflammatory gene expression during first 72 hours of response to cone ablation in *irf8* mutants shows minor differences compared to *irf8* heterozygotes

Since inflammation and microglia are implicated in MG-derived regenerative responses [14, 17, 20, 21, 26–28] and given the leukocyte/microglia responses seen in both *irf8* heterozygote and mutant (Fig 4), we examined expression of selected inflammatory genes at 24, 48, and 72 hours post treatment of *gnat2*:nfsb-mCherry larvae using RT-qPCR (Fig 6). For this analysis, we isolated RNA and prepared cDNA from pairs of eyes dissected from individual larvae treated with either DMSO or Mtz. We selected *il1b*, *il6*, *tnfa*, *tnfb*, and *il10* for the qPCR analysis because changes in these transcripts have previously been shown to be associated with retinal regeneration in zebrafish [16, 21, 50]. Although TNFα has been implicated in retinal regeneration in zebrafish [16, 51], we could not reliably detect *tnfa* with the primer pairs used in our study, therefore we could not assess *tnfa* mRNA changes. Although we detected *il10*, the Ct readings returned were highly variable and in high ranges (*il10* Cts ranged 30-36, we ran a total n=60 per genotype from both DMSO and Mtz treatments at each timepoint), indicating that detection of *il10* transcripts in larval retinal homogenates was not reliable nor comparable between samples; therefore, we did not complete analysis of *il10*. Because comparisons in the gene expression analysis involve two genotypes (*irf8*+/− and *irf8*−/−), two treatments (DMSO and Mtz), and three timepoints (24, 48, and 72 hpt), we provide the full results of statistical analysis in Supplemental File 1 and describe the prominent features of expression changes for each gene below. Graphs showing Log2FC over time are shown in Figure 6.

**Figure 6:**
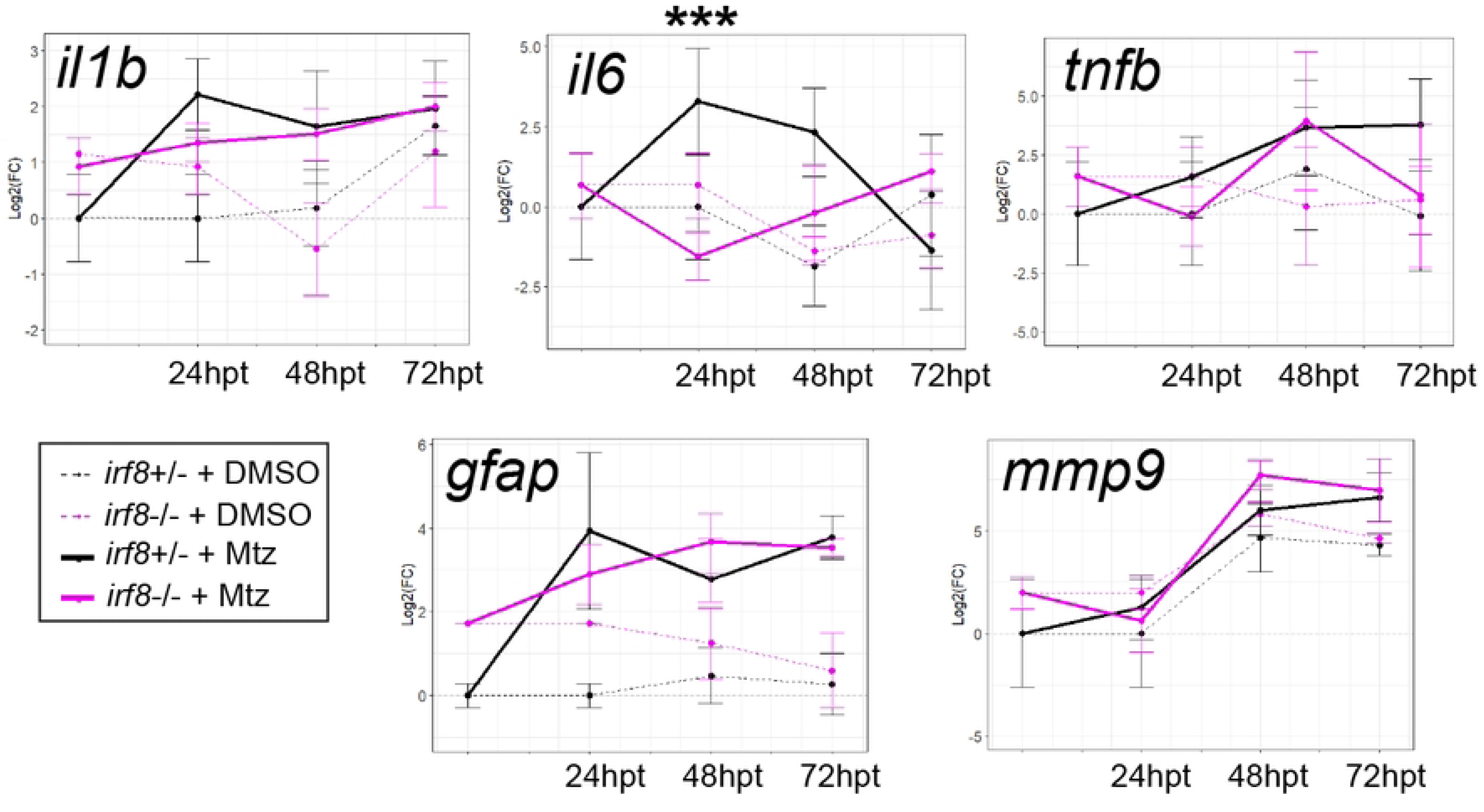
Time course of inflammatory gene expression and Müller glia reactivity genes following cone ablation in *irf8* heterozygotes and *irf8* mutants through 72 hours post treatment. RT-qPCR was performed using RNA collected at the indicated timepoints from pairs of whole larval eyes from *gnat2*:nfsb-mCherry larvae of either *irf8*+/− or *irf8*−/− genotypes, following DMSO or Mtz treatment. Gene expression for the indicated genes was analyzed compared to undamaged (DMSO treated) *irf8*+/− samples at 24 hpt, using the 2^ddCt method and graphing as Log2(FC). Complete results from statistical analysis are shown for this data in Supplemental File 1 and are discussed in the main text. For clarity, only one statistically significant result is annotated in this figure (***p=0.0002, 3-way ANOVA with Tukey’s posthoc) indicating statistically significant difference between *irf8*+/− and *irf8*−/− for *il6* expression following Mtz treatment.

Statistical results indicated that there were effects of timepoint, treatment (DMSO vs Mtz), and timepoint/genotype/treatment together on changes to *il1b* expression (Supplemental File 1). In general, *il1b* increased in *irf8*+/− over time following Mtz treatment, but levels of *il1b* also increased in DMSO treated *irf8*+/− samples by 72 hpt, indicating developmental changes in *il1b* mRNA also occur over this timeframe. In *irf8*−/−, *il1b* also increased but at an apparently more modest rate and showed increase in the DMSO group at 72 hpt. There were no statistically significant differences in *il1b* between *irf8*+/− and *irf8*−/− at any timepoints examined.

Statistical results indicated effects of all factors (timepoint, treatment, genotype), and all combinations of factors on *il6* expression (Supplemental File 1). In *irf8*+/−, *il6* increased by 24 hours of Mtz treatment compared to *if8*+/− treated with DMSO, remained elevated at 48 hpt, then decreased through 72 hpt. In *irf8*−/− samples, *il6* transcripts did not increase and instead remained similar to DMSO groups of both genotypes. The differences in *il6* for *irf8*−/− treated with Mtz compared to *irf8*+/− treated with Mtz at 24 hpt were statistically significant (Figure 6).

There were effects of timepoint and of treatment indicated for *tnfb* expression (Supplemental File 1). In general, *tnfb* levels showed a trend of increasing abundance in *irf8*+/−samples treated with Mtz over time, though differences were not statistically significant. The *irf8*−/− samples treated with Mtz showed a more variable trajectory with a statistically significant increase from 24 to 48 hpt (Figure 6 and Supplemental File 1). There were no significant changes in *tnfb* between *irf8*+/− and *irf8*−/− (Figure 6).

We also examined transcripts associated with Müller glia reactivity and damage responses by comparing expression of *gfap* [52, 53] and *mmp9* [21] in the two treatments and genotypes (Figure 6). Levels of *gfap* were initially increased in *irf8*−/− (DMSO) compared to *irf8*+/− (DMSO); although the differences were not statistically significant, they are consistent with our previous report comparing *gfap* between these genotypes [41]. In *irf8*+/−, *gfap* increased with Mtz treatment and remained elevated through 72 hpt. In *irf8*−/−, *gfap* also increased with Mtz treatment but perhaps at a more moderate rate (Figure 6). Overall, there were no statistically significant differences in *gfap* expression between *irf8*+/− and *irf8*−/− at specific timepoints.

The trajectory of *mmp9* expression in both *irf8*+/− and *irf8*−/− showed increases at 48 hpt and remained elevated at 72 hpt, and this was seen in both the Mtz treated groups as well as the DMSO treated groups compared to earlier timepoints (Figure 6). There were no statistically significant differences between genotypes at any specific timepoints. The results for *mmp9* indicated that increased expression in larval eyes over time is largely due to developmental changes, not an effect of Mtz treatment (Figure 6).

### Moderate reduction in early and late progenitor cell proliferation in *irf8* mutants following cone ablation

We next analyzed the amplification of MG-derived progenitor cell (MGPC) proliferation in *irf8*+/− and *irf8*−/− genotypes using PCNA staining and quantification in *gnat2*:nfsb-mCherry larval retina cryosections at several timepoints following cone ablation (Figure 7). Both genotypes showed increasing numbers of PCNA+ cells in the INL and ONL (consistent with MGPC proliferation and apical migration) through 120 hpt/5 days post treatment (dpt), followed by a reduction in numbers of PCNA+ cells at 144 hpt/6 dpt (Fig 7A-C). The peak of total PCNA+ cell numbers detected in retina cryosections was at 5 dpt for both *irf8* genotypes (Fig 7C). At 72 hpt/3 dpt, there were reduced numbers of PCNA+ cells in *irf8*−/− retinas compared to *irf8*+/− (Fig 7C), indicating a delay in the initial amplification of MGPC proliferation in microglia deficient conditions. Numbers of PCNA+ cells were moderately reduced for *irf8*−/− retinas at 96 hpt (4 dpt, but not statistically significant). Total numbers were similar between *irf8* genotypes at the peak detection of PCNA+ cell numbers at 120 hpt/5 dpt (Fig 7C), then *irf8*−/− retinas again showed reduced numbers of PCNA+ cells at 144 hpt/6 dpt (Fig 7C). When the numbers of PCNA+ cells were scored by distribution amongst retinal layers over the timepoints (Fig 7D), we found that the majority of PCNA+ cells were localized to the INL from 72-96hpt (3-4dpt) then at 120 hpt/5 dpt showed changes in distribution where PCNA+ cells were detected across both the INL and ONL (Fig 7D), consistent with the images in Fig 7A and with interkinetic nuclear migration described for MG-derived progenitor cells in regenerating zebrafish retinas [1]. The trends of PCNA+ cell distributions were similar between both *irf8*+/− and *irf8*−/− genotypes, but at 72 hpt/3 dpt there were fewer PCNA+ cells in the INL of *irf8*−/− retinas compared to *irf8*+/− (Fig 7D), again consistent with a delay in amplification of MGPC proliferation at this timepoint. Further, at 144 hpt/6 dpt, the *irf8*−/− retinas had fewer PCNA+ cells in the ONL than detected in *irf8*+/ (Fig 7D), consistent with reduced MGPC proliferation in later stages of the regenerative response.

**Figure 7:**
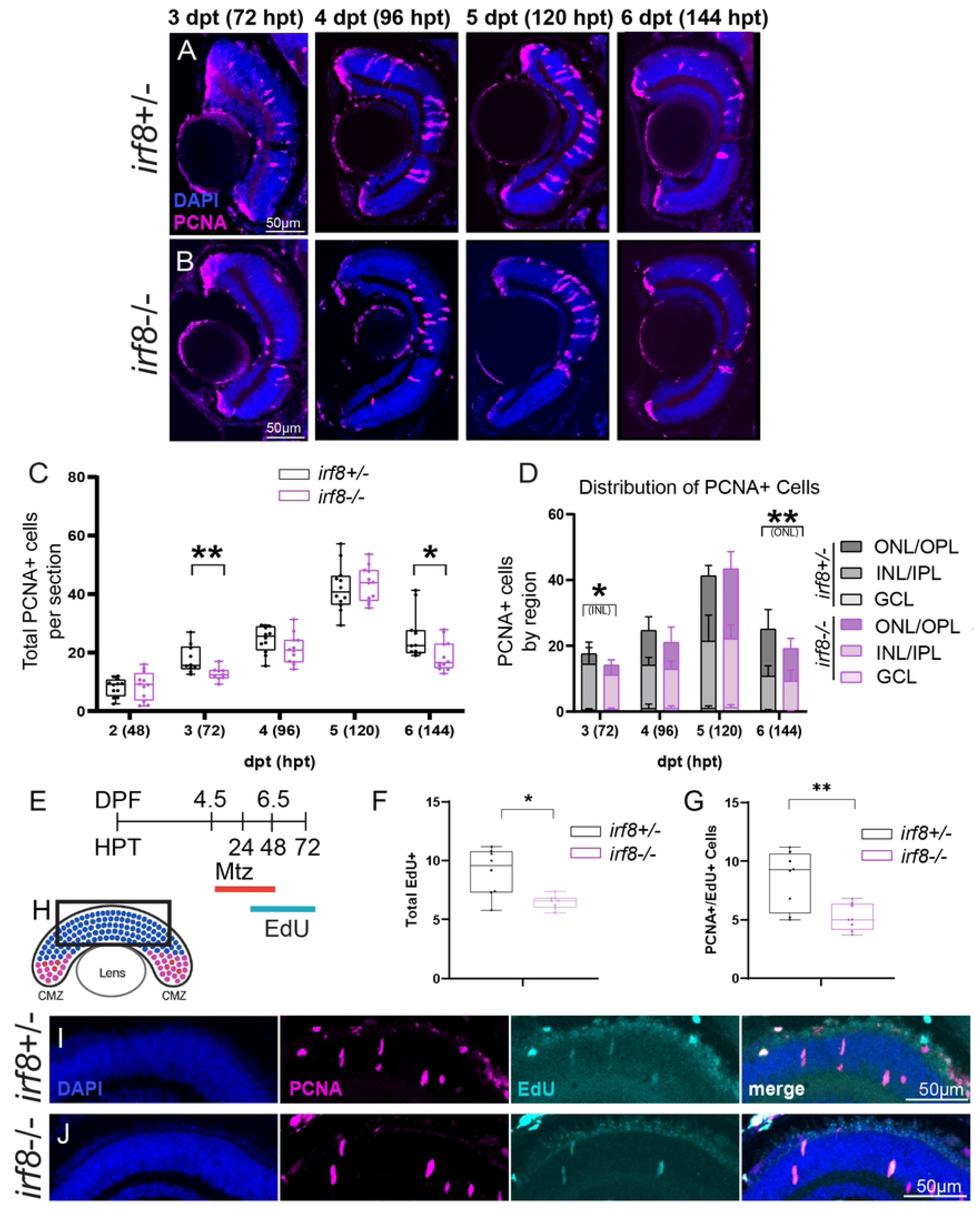
Amplification of MGPC proliferation over time shows minor differences in early and late cell cycle progression between *irf8* heterozygotes and mutants. A, B. Retinal cryosections from *gnat2*:nfsb-mCherry;*irf8*+/− or *gnat2*:nfsb-mCherry;*irf8*−/− larval zebrafish samples collected at the indicated timepoints post Mtz treatment and stained for PCNA and DAPI. Representative images are shown, scale bars in A and B apply to images in the respective row. C. Quantification of the total number of PCNA+ cells per retinal section in the two genotypes at the indicated timepoints. (The peripheral CMZ was excluded from this analysis). 2-way ANOVA (p<0.05) with post-hoc; annotation of all post hoc pairwise statistical comparisons on the graph were not included to prevent visual clutter. Pairwise comparisons indicated statistically significant increases in PCNA counts from 48 hpt (2 dpt) to 120 hpt (5 dpt) over time for both *irf8*+/− and *irf8*−/− with p<0.05 for comparisons across timepoints within the same genotypes, and for decreases in PCNA counts from 120 hpt (5 dpt) to 144 hpt (6 dpt) for both *irf8*+/− and *irf8*−/−. Annotated **p=0.0096, *irf8*+/− vs *irf8*−/− at 72 hpt (3 dpt) and *p=0.0189, *irf8*+/− vs *irf8*−/− at 144 hpt (6 dpt). D. Distribution of PCNA+ cells in the two genotypes by retinal layer at the indicated timepoints (ONL=outer nuclear layer, INL=inner nuclear layer, OPL=outer plexiform layer, IPL=inner plexiform layer). 3-way ANOVA (p<0.05) with posthoc, *p=0.0225 for INL counts *irf8*+/− vs *irf8*−/− at 72 hpt, **p=0.0028 for ONL counts *irf8*+/− vs *irf8*−/− at 144 hpt. E. Timeline showing treatment of *gnat2*:nfsb-mCherry;*irf8*+/− or *gnat2*:nfsb-mCherry;*irf8*−/− fish with Mtz and exposure to EdU. Quantifications of (F) the number of EdU+ cells (*p=0.0112, Mann Whitney U-test) and (G) the number of PCNA+EdU+ cells (**p=0.0076, Mann Whitney U-test) at 72 hpt in the two genotypes. H-J. Regions of retinal sections showing signal for DAPI, PCNA, and EdU relevant to quantifications in F and G. Scale bars apply to all images in the respective rows (I, J).

To further examine early MGPC proliferation at 72 hpt, we repeated previous experiments with EdU incorporation but extended EdU immersion exposure through 72 hpt/3 dpt (Fig 7E). We found reduced numbers of total EdU+ cells and PCNA+EdU+ co-labeled cells in *irf8*−/− retinas compared to *irf8*+/− retinas at 72 hpt/3 dpt (Fig 7F-J). These results, along with those for EdU incorporation at 48 hpt (Fig 5) and PCNA+ counts (Fig 7C), suggest that although entry into the cell cycle by responding MG is similar, MGPCs in *irf8*−/− may take longer to complete cell cycle progression than those in *irf8*+/− retinas during the early response to cone ablation. However, by 96 hpt, this delay has likely been mitigated and both genotypes show a similar peak of PCNA amplification at 120 hpt (Fig 7C). The reduction of PCNA+ numbers at 144 hpt/6 dpt indicates that MG-derived progenitor proliferation is trending down at this timepoint in both genotypes, however this may happen more rapidly in *irf8*−/− retinas and further reduce the proliferating MGPCs in the ONL (Fig 7C,D). Consistent with a reduction in MGPC proliferation between 120 and 144 hpt, our previous images suggested that regenerated cones may be present by these timepoints (Fig 2D, mCherry+ cells visible in the ONL). We therefore examined 120 hpt/5 dpt retinal cryosections for EdU+mCherry+ cones from experiments with EdU washout at 72 hpt. By this timepoint, EdU signal was considerably reduced, consistent with rapid proliferation of MGPCs from after EdU washout at 72 hpt, yet we were able to detect a few (on average, fewer than 3) EdU+mCherry+ cones in the central ONL of both genotypes at 120 hpt/5dpt (Supplemental Figure 2). Collectively, these results indicate that microglia may support cell cycle progression of MGPCs though the deficiency in *irf8* mutants results in only minor reductions in numbers of dividing progenitors at early and late stages of MGPC amplification. In addition, our results indicate that regenerated cones can be detected by 120 hpt/5 dpt in this experimental system.

We attempted to measure changes in re-expression of *ascl1a* by neuronal progenitor cells generated upon cone damage, using RT-qPCR (Supplemental Figure 3, statistical results shown in Supplemental File 1). There was a modest increase in *ascl1a* in *irf8*+/− detected at 24 hpt upon Mtz treatment, while increase in *ascl1a* was instead delayed to 48 hpt in *irf8*−/− upon Mtz treatment (Supplemental Fig 3). The active differentiation of neurons during development of the larval zebrafish retina may preclude the ability to detect significant changes in *ascl1a* and may be the reason we did not detect more robust changes in this gene upon Mtz treatment. However, these results are consistent with a delay in cell cycle progression in *irf8* mutants from 48 to 72 hpt (Fig 7C), in which the re-expression of *ascl1a* in MG-derived progenitor cells could also be temporally delayed.

### Regeneration of cones in larval retina is similar between *irf8* heterozygotes and mutants

We next analyzed regeneration of cones in both *gnat2*:nfsb-mCherry;*irf8*+/− and *gnat2*:nfsb-mCherry;*irf8*−/− larval retinas. After cone ablation, recovery of viable and healthy larvae later than 7 dpt (∼11 dpf age) was challenging with standard larval fish rearing conditions, which we attribute to the cone-dominant function of the larval zebrafish retina through approximately 15 dpf [54–56]. The loss of cones from Mtz-mediated ablation likely impacts larval feeding, and therefore survival; standard rearing of the DMSO treated larvae did not impact viability. Since we detected a small number of EdU+ regenerated cones at 5 dpt as described previously (Supplemental Fig 3), we selected 7 dpt as the first timepoint to analyze regenerated cones in the two *irf8* genotypes. To increase the likelihood of encountering food following cone ablation, larvae were maintained in glass beakers and provided ample powder food daily with a window of time to allow feeding before water/solution changes. All unhealthy or nonviable larvae were removed prior to powder food addition. In these experiments, we included EdU exposure by immersion of larvae in water including EdU from 96 hpt/4 dpt to 144 hpt/6 dpt, which corresponded to timeframes covering the peak of PCNA detection (Fig 7C).

At 7 dpt following Mtz treatment, mCherry+ cones were detectable in the ONL of both genotypes; several of these cones were EdU+ and centrally localized (and therefore not produced by growth at the CMZ) confirming that they were regenerated from MGPC (Fig 8A). Regions of the ONL in retinas from both genotypes still lacked cones and many of the mCherry+ cells within the central retina did not yet display typical cone morphology or patterns (Fig 8A), indicating that cone regeneration, differentiation, and/or maturation is ongoing at 7 dpt. We also included a stain for post-synaptic bipolar (BP) cell partners by staining for PKCα, which labels a subset of ON-BP types in zebrafish retina [57, 58] (Fig 8A). Patterns of PKCα+ signal showed BPs contacting cones in the ONL of both genotypes and also revealed that locations in which cones were absent had displaced BP cell bodies that protruded apically into the ONL region (Fig 8A). Further, some BP dendrites located basally to these “gaps” in cone signal appeared to “reach” for nearby cones tangential to the radial direction of the retina (Fig 8B); this was observed in both *irf8* genotypes. These results indicate that in the absence of pre-synaptic partners, BP cell bodies and dendrites remodel and reposition at least temporarily, during regeneration [44, 59, 60].

**Figure 8:**
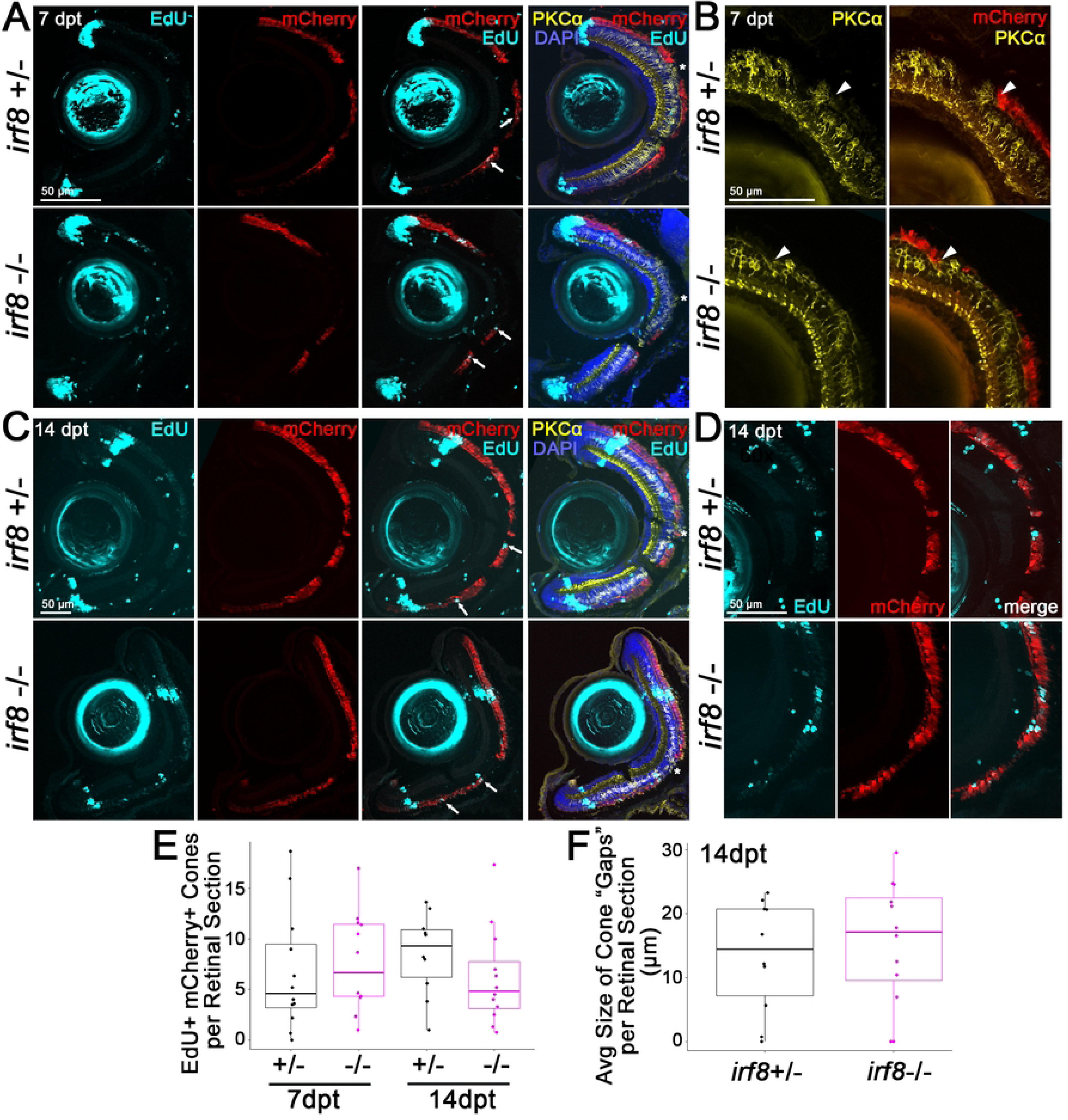
Regeneration of cones is similar for *irf8* heterozygotes and mutants. Retinal cryosections from *gnat2*:nfsb-mCherry;*irf8*+/− or *gnat2*:nfsb-mCherry;*irf8*−/− larval zebrafish were collected at 7- or 14-days post Mtz treatment (dpt), with EdU exposure from 4-6 dpt. After EdU retrieval, cryosections were stained for PKCα and DAPI, mCherry signal was used to visualize cones. Selected images and individual or merged channels are shown for retinal sections from the two *irf8* genotypes collected at 7 dpt (A) and at 14 dpt (B). Arrows annotate EdU+ mCherry+ cones, * indicates regions where PKCα+ bipolar (BP) cell bodies are displaced into the ONL. C. Images show PKCα+ BP cells with dendrites extended and contacting cones at locations adjacent to the radial direction of the retina in samples from both *irf8* genotypes at 7 dpt. D. Images are higher magnification views of EdU+ mCherry+ cones in retinal cryosections collected at 14 dpt; several EdU+ mCherry+ cones are detected in the central retina of both *irf8* genotypes as well as other EdU+ cells in the inner retina. E. EdU+ mCherry+ cells (cones) were quantified per retinal section in the two genotypes at both timepoints (the peripheral CMZ was excluded from this analysis and counting was performed inside the “stripes” of EdU seen in the 14 dpt samples). Differences between genotypes and between timepoints were not statistically significant (Mann-Whitney U-test for pairwise comparisons between *irf8*+/− and *irf8*−/− at 7 dpt and at 14 dpt, and pairwise comparisons within the same genotype between 7 dpt and 14 dpt). F. Quantification of the size of gaps in mCherry+ signal in the ONL at 14 dpt; differences were not statistically significant (Mann-Whitney U-test).

We also performed experiments with similar timing of EdU exposure by immersion, with collection at 14 dpt, to examine regeneration at a later timepoint (Figure 4C). For these experiments, larvae were again exposed to EdU in glass beakers with feeding of ample powder food. Due to challenges with survival to this experimental endpoint and to ensure we could collect enough larvae, some experiments with collection at 14 dpt used multiple breeder pairs and clutches of larvae with known *irf8* genotypes. We did not find differences in eye size measurements between *irf8*+/− and *irf8*−/− samples at 14 dpt (Supplemental Fig 4). After EdU washout, larvae were transferred to a specialized housing apparatus in fish tanks and provided close feeding with powder and brine shrimp until tissue collection (details described in Methods). At 14 dpt, EdU+ cones were detected in regions of the ONL consistent with cone regeneration in samples from both genotypes: these cells were located interior of “stripes” of EdU visible in retinal sections, representing a boundary of growth from the CMZ formed during the EdU exposure window (Fig 8C,D). Quantification of EdU+mCherry+ cones at 7 dpt and at 14 dpt did not find statistically significant differences between *irf8*+/− and *irf8*−/− samples (Fig 8E).

We also noted that the number of EdU+mCherry+ cones did not increase in retinas collected at 14 dpt compared to those at 7 dpt (Fig 8E); however, the ONL appeared more populated with mCherry+ cones (Fig 8C). These observations indicate that regeneration is ongoing from 7 to 14 dpt, and not all regenerated cones are retained over time and/or EdU label is further diluted due to ongoing progenitor cell divisions. Where “gaps” in mCherry+ cone signal in the ONL were observed at 14 dpt, we again observed apically displaced BP cell bodies (Fig 8C). We measured the curvilinear distance of the “gaps” in mCherry+ cone signal in the ONL and did not find differences in gap sizes between *irf8* heterozygotes and *irf8* mutants (Fig 8F), again indicating that regeneration was occurring similarly between the two genotypes.

Some EdU+ cells were also detected in the inner retina at both 7 and 14 dpt that were not mCherry+ cones (Fig 8A,C,D), indicating that our EdU immersion schedule labeled other cell types. These could represent MG that were in the cell cycle at the time of EdU exposure and/or other neurons that were produced during this timeframe, as production of neuronal cell types different than the ones targeted has been described for zebrafish retinal regeneration [61–63]. The number of EdU+ cells that were scored as mCherry-negative “non-cones” were not statistically different between *irf8* genotypes (Supplemental Fig 4).

To examine potential synapse formation of regenerated cones with BPs, we visualized OPL synaptic regions with higher magnification, and qualitatively compared features visualized in undamaged retinal cryosections (Figure 9A,C) to samples collected at 7- and 14-days post Mtz treatment, from both *irf8* genotypes (Figure 9B,D). Staining for SV-2 (Synaptic Vesicle 2) was included in these images to visualize pre-synaptic vesicle+ regions in mCherry+ cones in relation to cone-BP contacts. SV-2 signal was detectable in the OPL, showing localization consistent with that of cone pedicles, of both *irf8* genotypes at 7 and 14 dpt following cone ablation, though it was less uniform and appeared “patchier” in samples that had experienced cone ablation compared to undamaged retina (Fig 9B,D). In samples with regenerated cones, cone morphology did not fully recapitulate that of undamaged cones (upright in the radial direction of the retinal section, with clear axons and pedicles). Instead, cones in retinas following ablation were less elongated with axons harder to discern, often obliquely oriented relative to the radial axis of the tissue section, appeared disorganized, and this was the case for both *irf8* genotypes (Fig 9B,D). As noted above, BP cell bodies were seen protruding into the ONL where gaps in cone signal were present (Fig 9B,D). However, regions with features consistent with re-established synapses (mCherry+ cone basal pedicle contacting PKCα+ BP dendrites also co-labeling with SV-2) were notable in samples collected at 7 and 14 dpt following Mtz treatment (Fig 9B,D,E,F). Visualizing these features in the ONL selectively for regions with EdU+ mCherry+ cones, confirmed that SV-2+ synapses were restored between regenerated cones and BPs and this was evident for both *irf8* heterozygotes and mutants (Fig 9E,F).

**Figure 9:**
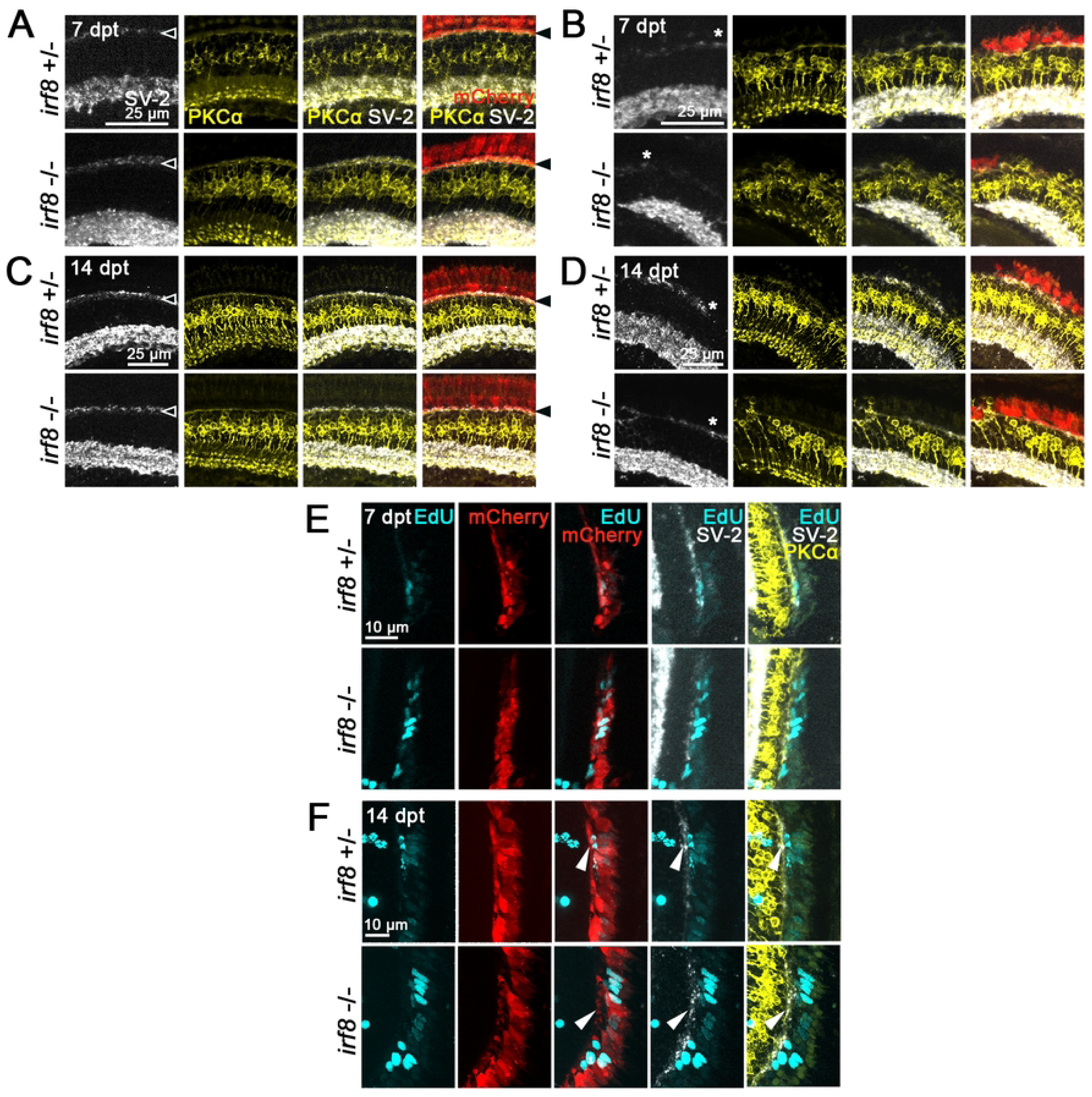
Synapses in the OPL are evident for regenerated cones in both *irf8* heterozygotes and mutants. Retinal cryosections from *gnat2*:nfsb-mCherry;*irf8*+/− or *gnat2*:nfsb-mCherry;*irf8*−/− larval zebrafish were collected at 7 or 14 days post DMSO or Mtz treatment (dpt) and included EdU exposure as described in the text. Cryosections were stained for PKCα, SV-2, and DAPI; mCherry signal was used to visualize cones. A. Shows intact and typical outer retinal morphology in cryosections from undamaged samples (DMSO treated) collected at 7 dpt, from each *irf8* genotype. Outer plexiform layer (OPL) is visualized by SV-2 signal (white arrowhead in left panel) where cones and PKCα+ bipolar cells have synapses (black arrowhead, right). B. Shows images from selected samples collected at 7 dpt following Mtz treatment from each *irf8* genotype. SV-2 signal is dim and patchy (annotated by *) though cones and BPs are again showing contact in the regenerating OPL. C. Shows intact and typical outer retinal morphology in cryosections from undamaged samples (DMSO treated) collected at 14 dpt, from each *irf8* genotype. Outer plexiform layer (OPL) is visualized by SV-2 signal (black arrowhead, right) where cones and PKCα+ bipolar cells have synapses. D. Shows images from selected samples collected at 14 dpt following Mtz treatment from each *irf8* genotype. SV-2 signal is more apparent and stronger but still patchy (annotated by *) though cones and BPs are again showing contact in the regenerating OPL. Histological abnormalities (as described for Fig 8) are also visible here where BPs protrude into the OPL. E, F. Shows enlarged regions of images acquired from samples with cone regeneration where EdU+ (regenerated) mCherry+ cones have re-established contacts with PKCα+ bipolar cells. Samples were collected at 7 dpt (E) and at 14 dpt (F). The white arrows point out SV-2+ localization where EdU+ regenerated cones contact PKCα+ bipolar cells in the OPL.

### Compensatory immune cell populations in *irf8* mutants with differential staining of 4C4 and Lcp1 markers are present during regeneration of cones

Given the overall lack of significant differences observed in regenerative responses in *irf8* mutants compared to heterozygotes and given the detection of some leukocytes/microglia at 48 hpt/2 dpt in *irf8* mutant retinas (Figure 4), we performed further analysis of microglia/leukocytes at additional, selected timepoints following ablation of cones in *gnat2*:nfsb-mCherry larvae of both *irf8* genotypes (Fig 10). For this analysis, we again stained retinal cryosections with the 4C4 antibody [45] and the pan-leukocyte marker Lcp1 [47]. We performed this analysis at timepoints corresponding to ablation of cones (48 hpt, Fig 1D, Fig 4), surrounding the peak of PCNA+ MGPC numbers (5-7 dpt, Fig 7C), and timepoints for which we examined cone regeneration (7 and 14 dpt, Figures 8 and 9). Comparing total numbers of cells staining with these markers (cells that stained positive for either one marker or both markers), we found that in retinas of *irf8* heterozygotes, immune cell populations expanded at 5 dpt compared to 48 hpt/2 dpt, then declined from 6 dpt through 14 dpt (Fig 10A). In comparison, *irf8* mutant retinas had reduced numbers of immune cells at all timepoints compared to *irf8* heterozygotes, though they did have detectable immune cell populations (Figure 10A). The immune cell populations in *irf8* mutant retinas also showed expansion at 5 dpt and contraction by 14 dpt (Figure 10A). At 14 dpt, on average less than 2 immune cells were detected per retinal cryosection in *irf8* mutants (Fig 10A). The populations detected in *irf8*+/− retinas at each of these timepoints nearly all stained positive for Lcp1, with the majority also co-labeling with 4C4 (Figure 10B-E). Interestingly, within the populations of immune cells detected in *irf8* mutants from 5 to 7 dpt was a subpopulation of cells that stained positive only for the 4C4 marker, and not Lcp1 (Figure 10B-E). This “4C4-only” population localized primarily to the ONL of *irf8* mutant retinas from 5-7 dpt but at 14 dpt was also present in the inner retina of *irf8* mutants, but not *irf8*+/− (Figure 10D,E). This analysis revealed that immune populations are reduced in numbers in *irf8* mutants compared to *irf8* heterozygotes, but compensatory immune cell populations are present during regeneration of cones. The differential labeling with 4C4 antibody suggests that these compensatory populations in *irf8* mutants are phenotypically different than those in *irf8* heterozygotes.

**Figure 10:**
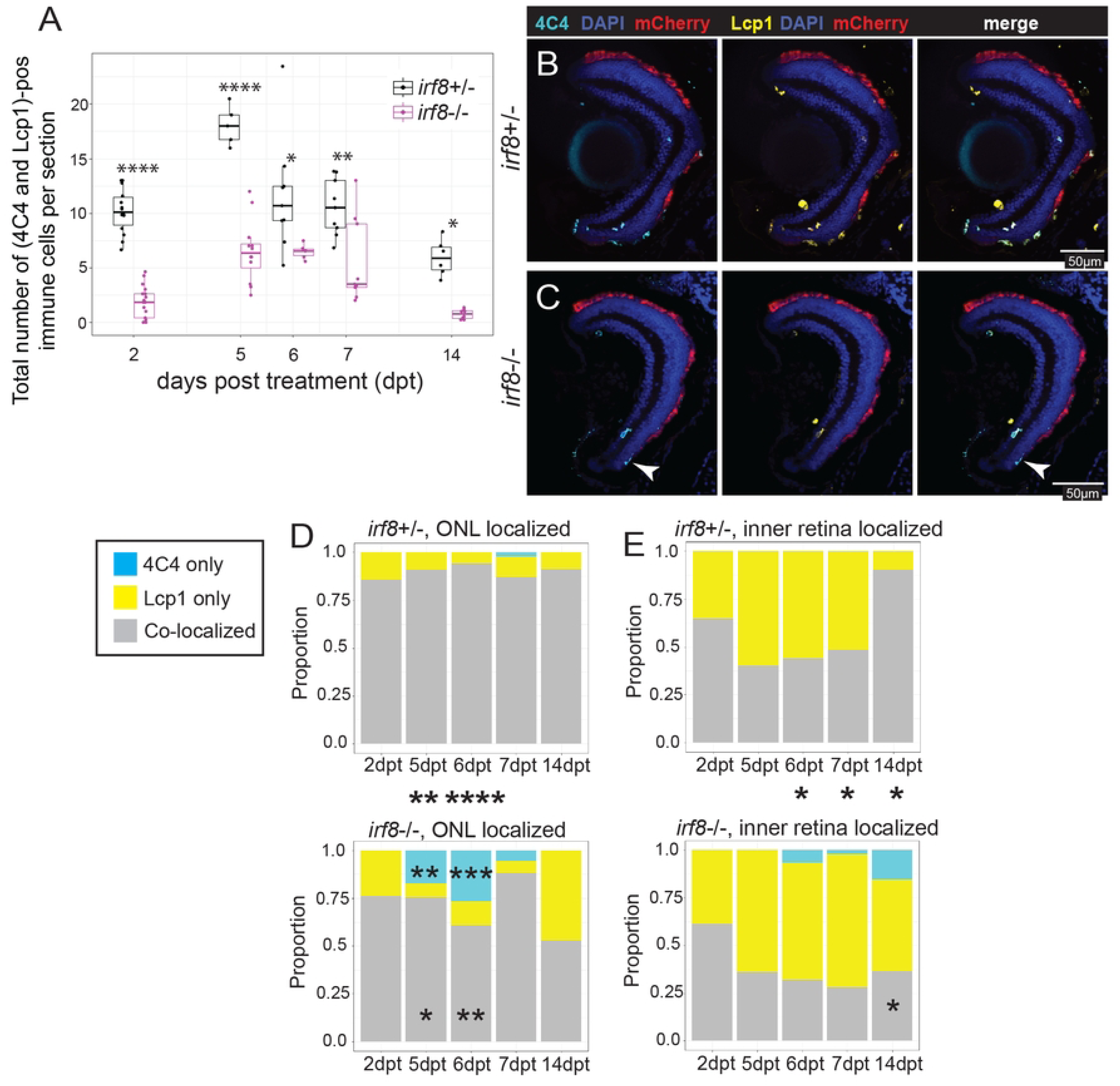
Analysis of immune cell numbers and marker expression in *irf8* heterozygotes and *irf8* mutants over time. A. Quantification of the total number of 4C4+ and L-plastin (Lcp1+) cells (cells counted here included those that stained positive for one or both markers) in retinal cryosections of *gnat2*:nfsb-mCherry;*irf8*+/− or *gnat2*:nfsb-mCherry;*irf8*−/− larval zebrafish collected at the indicated timepoints post Mtz treatment. 2-way ANOVA (p<0.05) with Tukey’s posthoc. P-values are annotated only for pairwise comparisons between *irf8*+/− and *irf8*−/− (*p<0.05, **p<0.01, ***p<0.001, ****p<0.0001). All p-values between counts at each timepoint compared to the others were <0.05 for both genotypes. B. The selected images shown here are from 7 dpt samples. Images of retinal cryosections stained for DAPI, 4C4, and Lcp1; cones are visualized by mCherry fluorescence. Images show populations that co-label for both 4C4 and Lcp1 (all cells in the ONL of *irf8*+/− images here) as well as cells that are 4C4+ but do not have Lcp1 signal detected in the outer nuclear layer of *irf8* mutants (arrowheads). D, E. Proportions of immune cells staining for the 4C4 and Lcp1 markers, localized to the ONL (D) or to the inner retina (E), in the two *irf8* genotypes at the indicated timepoints. P-values <0.05 (Fisher Exact test) are annotated at selected timepoints where there was a statistically significant difference in *irf8*−/− compared to *irf8*+/−; *p<0.05, **p<0.01, ***p<0.001. Asterisks between the two graphs indicate differences between the distributions at the indicated timepoints for *irf8*−/− compared to *irf8*+/−. Asterisks within the *irf8*−/− graph indicate where there were different proportions of cells with the indicated markers *irf8*−/− compared to *irf8*+/−.

## Discussion

We examined the regenerative response in the larval zebrafish retina following ablation of Nfsb-expressing cone photoreceptors achieved by exposure to metronidazole. Experiments were performed in microglia-sufficient larvae and in microglia-deficient *irf8* mutants, using siblings wherever possible in our experimental design. We performed analysis over a time course spanning 14 days post-cone ablation and examined the ablation of cones, early glial cell responses, the first asymmetric division of the MG, amplification of MGPC proliferation, expression of selected inflammation-associated genes, and the regeneration of cones and their synapses with BPs. We also examined microglia and other immune cells (by staining with 4C4 and Lcp1 antibodies) at multiple timepoints in both *irf8* heterozygotes and *irf8* mutants. We found that overall, regeneration proceeds sufficiently in *irf8* mutant larval zebrafish with moderate reduction in early and late MGPC proliferation. Notably, though *irf8* mutant zebrafish retinas are highly deficient in microglia/immune cells prior to induction of damage, we identified a population of immune cells that emerge in *irf8* mutant retinas following cone ablation. This compensatory immune cell response precludes the ability to make strong conclusions about the role of microglia in retinal regeneration in zebrafish based on our experiments presented here. Nonetheless, our results are consistent with multiple reports [14, 15, 31, 41] which, when considered together with our work presented here, collectively indicate that (i) microglia likely support the proliferation of MGPC in the zebrafish retina following damage and (ii) that compensatory mechanisms and cell types provide function in microglia-deficient retinas.

Other studies utilizing the *irf8*^st95^ [40] mutant zebrafish line to manipulate microglial abundance during retinal regeneration have been published previously [15, 18, 36] and two of these reports involved photoreceptor regeneration (MG-mediated) in adult fish [15, 18]. Bludau et al. [15] found a modest decrease in proliferating progenitors in the damaged *irf8* mutant retina following light damage to photoreceptors, while Song et al. [18] found no difference in proliferating cells in the retina in *irf8* mutants following light damage. These reports appear conflicting; however, it is worth noting that only one timepoint was examined in analysis of progenitor proliferation in these two studies, and the selected timepoint was a day apart. The other study using the *irf8* mutant, by Leach et al. [36], examined regeneration of the RPE in larvae and is not known to involve MG-mediated progenitors towards the regeneration of RPE cells. Here, we performed analysis of MGPC proliferation in *irf8* mutants and heterozygous siblings following cone photoreceptor damage over a more comprehensive time course, as well as the initial entry of responding MG into the cell cycle. We found that in *irf8* mutants, MG entered the cell cycle with similar timing (by 48 hours post drug exposure); however, early (72 hpt) and late (144 hpt) MGPC proliferation were reduced compared to sibling controls. Our results are consistent with findings for MGPC proliferation during retinal regeneration in larval zebrafish following rod ablation which manipulated microglia using a different strategy [14].

Further, multiple reports using dexamethasone to suppress microglia and inflammatory signals found impacts on MGPC proliferation [14, 15, 21, 22]. Collectively, the emerging theme is that inflammatory signals promote MGPC proliferation in the damaged zebrafish retina and microglia are a likely source of such signals. Consistent with this, we found moderate changes in inflammatory gene expression in *irf8* mutants following cone ablation, particularly for the cytokine *il-6*. A study using Dexamethasone and *irf8* mutant zebrafish in light damaged retinas [32] supports the idea that microglia promote some inflammatory component of the response to retinal damage, which acutely may be important for the MGPC response but that could be detrimental in contexts of chronic photoreceptor degeneration. The molecular nature of such signals remains to be fully identified, and it is likely that multiple cell types can provide such signals. Moreover, microglia may function in an inter-cellular communication axis involving not only the MG but other cell types such as vascular endothelial cells [31].

Our analysis of regenerated cones did not detect differences in *irf8* mutants compared to their siblings. Regenerated cones were detected at similar abundance in both genotypes, which we observed to make synaptic connections with BPs in the regenerating OPL. This is in contrast to previous studies that found delayed photoreceptor regeneration in laser damaged retinas when microglia were depleted with PLX3397 treatment [20], and that found delayed/reduced replacement of rods in larval retina when microglia were depleted using an Nfsb/Mtz strategy [14]. In addition, the regeneration of RPE in larval zebrafish eyes was reduced/abnormal when microglia were depleted using genetic and pharmacological tools [36, 43]. The different outcome for photoreceptor regeneration in our studies here (which was overall, normal) could be the age of fish used in the experiments, the presence of compensatory immune cell populations in *irf8* mutants (discussed below), the methods used to quantify regenerated photoreceptors/RPE cells, the timepoints selected for analysis, stem cell niches that do not involve MG and are available for regenerated RPE cells [64, 65] and for rods [2], or other factors. Further, though we qualitatively described detection of synaptic connections of regenerated cones with BPs in the OPL in retinal cryosections, we did not deeply or quantitatively analyze regenerated synapses. It remains unknown whether microglia modulate synaptic rewiring in regenerated retinas [7, 44].

As in previous reports using *irf8* mutant zebrafish [15, 18, 36], we detected immune cells in *irf8*−/− retinas following neuronal damage. In our studies described in this paper, these other leukocytes expanded upon retinal damage then were reduced over time, with a trajectory similar to their microglia-sufficient siblings but at markedly reduced abundance. Interestingly, the leukocyte populations detected in *irf8* mutants in our experiments stained positive for markers L-plastin (Lcp1) and 4C4 but did not always express both markers. Notably, we detected a subset in microglia-deficient *irf8* mutants that stained positive with 4C4 antibody but negative for Lcp1 and showed increased abundance compared to the microglia-sufficient siblings. This could indicate different phenotypes and/or ontogeny of the immune cells evoked in *irf8* mutant retinas compared to microglia-sufficient retinas. The compensatory immune cell response suggests an essential role for the immune system in retinal damage and possibly for regeneration. The ability of non-hematopoietic cell types, such as the neuroglia, to execute compensatory functions has been described [41, 66, 67]. Such responses may be limited and/or scale with the extent and nature of damage to retinal neurons.

The presence of such compensatory immune cell populations, however, makes it difficult to draw conclusive interpretations from our experiments in this paper. This has been emphasized previously [18] and more recently, others have instead used zebrafish with loss of function mutations in both *csfr1a* and *csfr1b* [31, 43] to manipulate microglial abundance. Further, drugs such as PLX3397 which blocks CSFR1 signaling have been used to manipulate microglia in zebrafish [20, 36]. However, in our results PLX3397 treated retinas had even more leukocytes than *irf8* mutants, therefore we elected to forgo this approach. Though the use of *irf8* mutant zebrafish may not be ideal for some experiments, such as those involving retinal damage that elicit compensatory immune cell responses as we and others found, this microglia-deficient *irf8* mutant still has a place for developmental studies probing microglial functions. Indeed, we report here that undamaged larval *irf8* mutant retinas are strongly deficient in microglia, consistent with our previous work [41]. The *irf8* mutant offers advantages over the need for extensive genetic crosses to generate *csfr1a/b* double mutants and for prolonged drug exposures aimed to deplete microglia. Nonetheless, caution should be used in interpreting experiments with any microglia-deficient system as compensatory responses from not only other immune cells but also neuroglia may be elicited in the absence of microglia [41, 66, 67].

Though our findings presented in this paper are limited and not fully conclusive, other literature emphasizes the impact of microglia on the outcome of retinal regeneration [27, 28, 31, 39, 43]. This is an important area of scientific pursuit given the involvement of microglia in numerous neurodegenerative diseases, many of which may involve compensatory cellular and molecular mechanisms and pathways. Continued interrogation of microglial functions will require focused, targeted approaches with advanced experimental design.

## Methods

### Animals

All experiments used zebrafish (Danio rerio) and procedures were approved by the University of Idaho Animal Care and Use Committee (IACUC). Adult zebrafish were housed in an aquatic system with filtered, recirculating water, maintained at 28°C and on a 14:10 light:dark cycle as described in Westerfield, 2007 [68]. Light onset was considered 0 hours post-fertilization (0 hpf). Upon fertilization, embryos were collected into glass beakers with system water and either reared in the fish facility or transferred to an incubator with 14:10 light:dark cycle at 28°C until treatments commenced. Water for embryos/larvae was refreshed daily. All experiments used transgenic *gnat2*:nfsb-mCherry fish [23] in which cone photoreceptors express bacterially derived nitroreductase tagged with mCherry under the cone-specific promoter *gnat2*. In some analyses, the *gnat2*:nfsb-mCherry line was crossed to the *TP1*:mTurquoise^uoi2523Tg^ transgenic line [29]. The *gnat2*:nfsb-mCherry fish were crossed with *irf8^st95* [40] homozygous mutant partners, to generate *gnat2*:nfsb-mCherry;*irf8*+/− breeders. The *gnat2*:nfsb-mCherry;*irf8*+/−breeders were then crossed to *irf8* mutants (*irf8−/−)* to generate *gnat2*:nfsb-mCherry+ microglia-sufficient (*irf8*+/−) and -deficient (*irf8*−/−) offspring for experiments. Whenever possible, experiments were performed blind to genotype at the treatment and collection stages, using clutches of offspring with unknown *irf8* genotype. Genotyping for the *irf8^st95^* mutation was performed after sample collection; gDNA extraction and the PCR/AvaI restriction digest protocol as previously described [40, 41]. Feeding of larvae began at 6 dpf with powder food provided once daily. At 10 days post-fertilization (dpf), larvae were transferred to feeding compartments in tanks and received brine shrimp in the morning and powder food in the afternoon.

### Treatments

At 4 dpf, embryos were screened using a fluorescent sorting microscope for the presence of mCherry signal in eyes (also visible in pineal gland), indicative of transgene expression in cone photoreceptors. Solutions were prepared by dissolving dimethyl sulfoxide (DMSO, vehicle) to 0.1% final concentration with or without metronidazole (Mtz, Thermo Scientific 210341000) at 10 mM final concentration in fish water. Solutions of DMSO/Mtz were protected from light and mixed for a minimum of 4 hours on an orbital shaker before use. Embryos positive for mCherry were split into two groups (genotypes unknown) and immersed in either 0.1% DMSO or 10 mM Mtz for 48 hours total (approximately 4.5 – 6.5 dpf), with refreshment of treatment solutions at 24 hrs. Because Mtz is light sensitive, these treatments were done in petri dishes covered loosely with foil. After treatments, larvae were removed from the dishes and transferred to beakers with fresh water for washout and recovery. In some experiments, larvae were immersed in 175 μM 5-ethynyl 2’-deoxyuridine (EdU, Invitrogen) for 24-48 hours total, with EdU solutions refreshed at 24-hour intervals. Feeding of powder food was administered prior to the time of EdU solution refreshment.

For regeneration timepoints (collection at 7- or 14-days post treatment, dpt): at 10 dpf (6 dpt = 144 hpt), the larvae were transferred to a feeding compartment within a 2.8L tank. The feeding compartment uses a 0.5L plastic container with a fine mesh screen bottom with edges that extend and rest on the lip of the tank, allowing the feeding compartment to sit under the feeding hole and water feed of the tank lid while on the aquatic housing system. The larvae were placed in the 0.5L feeding compartment, permitting food to be temporarily restricted to the location of the larvae, increasing chances of successful feeding. The mesh bottom allows for constant water flow that slowly removed the excess brine shrimp and powder food from the compartment, maintaining water quality. Feeding larval zebrafish in this housing situation consisted of morning brine shrimp (pre-chilled at 4°C to reduce locomotion of the shrimp) and afternoon powder feedings of ∼0.05g/feeding. Buildup of excess food was removed daily by gentle aspiration with a plastic transfer pipette to maintain water quality.

### PLX3397 treatments

Solutions were prepared by dissolving DMSO to 0.1% final concentration with or without PLX3397 (Sellick Chemicals) at 1µM final concentration in fish water. Solutions were protected from light during mixing and treatment. At 2 dpf, embryos were dechorionated and split into treatment groups. Embryos were treated with either DMSO or PLX3397/DMSO with solution refreshes occurring every 24 hours until 4.5 dpf, when they were sorted for expression of the *gnat2*:nfsb-mCherry transgene. Embryos positive for mCherry signal were split into 4 groups based on prior PLX3397 treatments: DMSO, PLX3397/DMSO, Mtz, and PLX3397/Mtz. Mtz solutions were prepared as previously described, either with or without the addition of 1µM PLX3397. Solutions were refreshed every 24 hours until collection at 6.5 dpf.

### Cryopreservation and sectioning of retinal tissue

On the day of sample collection, larvae were euthanized with 0.4% v/w tricaine/MS-222 and then fixed in 4% paraformaldehyde in 5% sucrose phosphate overnight at 4°C with constant rocking. The following day, larvae were washed in a graded series of phosphate buffered solution (pH 7.4) of 5% to 20% sucrose. Tails were clipped from fixed larvae and transferred to hot NaOH gDNA extraction buffer [69] then genotyped for *irf8^st95^* mutation as described above. Samples were transferred to a 1:2 solution of OCT embedding medium (Sakura Finetek): 20% sucrose phosphate and mixed for 30 minutes at room temperature. Larvae were embedded in 1:2 OCT:20% sucrose phosphate and frozen by immersion in 2-methylbutane supercooled with liquid nitrogen. After freezing, blocks containing larvae were stored at −20°C for at least 24 hours. Sectioning was performed using a Leica CM3050 cryostat, with 10 μM thick sections placed onto glass slides (FisherBrand Superfrost Plus Microscope Slides). Sections on slides were desiccated overnight then stored at −20°C until use.

### Immunostaining, TUNEL labeling, and EdU detection

Retinal cryosections on slides were thawed in a humidified chamber at room temperature. For PCNA staining, citrate buffer antigen retrieval was performed prior to the blocking step as previously described [13, 70]. TUNEL staining was performed following the manufacturer’s instructions (Roche), and slides were rinsed in PBS, prior to blocking and antibody staining. EdU detection, where appropriate, was completed using an EdU Click-iT kit (Invitrogen) or EdU Assay/EdU Staining Proliferation Kit (iFlour 647) (Abcam) following the manufacturer’s instructions. EdU detection was completed prior to immunostaining, then slides were transferred to antibody blocking buffer and stained as described below.

Blocking solution (20% normal donkey serum, 0.1% sodium azide, 0.1% Triton X-100, prepared in phosphate buffered saline (PBS)) was added to slides and incubated for 30 minutes at room temperature. Primary antibodies were prepared in antibody dilution buffer (1% normal donkey serum, 0.1% sodium azide, 0.1% Triton X-100, prepared in PBS) then added to slides and incubated overnight at 4°C. The following day, slides were washed in PBS + 0.1% Triton-X-100 (PBST) at room temperature for 30 minutes. Secondary antibodies and DAPI were prepared in antibody dilution buffer. Slides were incubated in secondary antibody solution at room temperature for at least 1 hour. Slides were washed for 30 minutes in PBST, rinsed in PBS, then mounted with coverslips using Vectashield Vibrance.

Primary antibodies and dilutions were as follows: mouse 4C4 antibody (1:100, used as hybridoma supernatant from clone 7.4.C4 sourced from Sigma-Aldrich 92092321), rabbit anti-Lplastin (Lcp1, 1:10,000, a kind gift of Dr. Michael Redd), rabbit anti-proliferating cell nuclear antigen (PCNA; 1:200, Proteintech 10205), mouse anti-Glutamine Synthetase (GS; 1:200, BD Biosciences 610517), mouse anti-SV2 (1:2000, Developmental Studies Hybridoma Bank), Rabbit anti-PKCa (1:500,Santa Cruz Biotechnology). Secondary antibodies were from Jackson Immunoresearch and used at 1:200 dilution (donkey anti-mouse and donkey anti-rabbit antibodies conjugated to Alexa-Fluor488, Cy3, or Alexa-Fluor647) or from AAT Bioquest (goat anti-mouse Cy7).

### Confocal microscopy, image processing, and analysis

Imaging was performed on a Nikon CrestOptics X-Light spinning disc confocal microscope running Nikon Elements software, using 20X dry (Plan Apo λ 20X Air 0.75 NA DIC), 40X water immersion (Apo LWD λS DIC N2 1.15 NA), or 60X oil immersion (Plan Apo λ 40X oil 1.40 NA WD=130µm) objectives. Z stacks (optical sections) were collected at 0.3-3 micron step sizes. Images were viewed and analyzed using ImageJ (FIJI) or Nikon Elements software. For retinal cryosections, 3-6 sections per eye were imaged (representing technical replicates) from which the average of technical replicates was determined for each sample that was quantified/counted. Sections containing central retina were selected for analysis. Counting of microglia, PCNA/EdU+ cells, and/or eye measurements were performed in entire retinal cryosections, using cell specific markers (from immunostaining or fluorescent reporters) and signal from the nuclear stain DAPI to identify retinal layers.

### RNA extraction and cDNA synthesis

On day of collection, larvae were euthanized with 0.4% w/v tricaine, water was removed, and the larvae were flash frozen on liquid nitrogen. Flash frozen larvae were stored in 100% methanol at −80°C until RNA extraction. Pairs of eyes were dissected from larval heads and transferred to RNA lysis buffer then immediately homogenized with a handheld homogenizer (Bio-Gen Series PRO200, Pro Scientific). RNA was extracted using Zymogen Quick-RNA microprep kit following the manufacturer’s instructions. After eye removal, bodies were transferred to gDNA extraction buffer followed by *irf8* genotyping. RNA was quantified and checked using a NanoDrop^TM^ One Microvolume UV-Vis Spectrometer (Thermo Scientific). Following RNA extraction, cDNA was synthesized using SuperScript IV First-Strand cDNA Synthesis Kit (Invitrogen) using random hexamers. The cDNA samples were stored at −20°C until qPCR.

### Quantitative PCR (qPCR)

Gene-specific primers for qPCR and their sources are shown in Table 1. Quantitative PCR (qPCR) was performed using PowerTrack SYBR-Green Master Mix (Applied Biosystems), using 2 ng of cDNA template per reaction, with reactions done in technical duplicates. Reactions were run on a QuantStudio^TM^ 3 Real-Time PCR machine (ThermoFisher). qPCR reactions underwent 40 amplification cycles, and non-template controls were included for all primer pairs. After reactions, amplification and melt-curves were inspected for clean, specific peaks. Fold change was calculated using the delta delta Ct method using *elf1a* Ct readings as the calibrator gene.

**Table 1.**
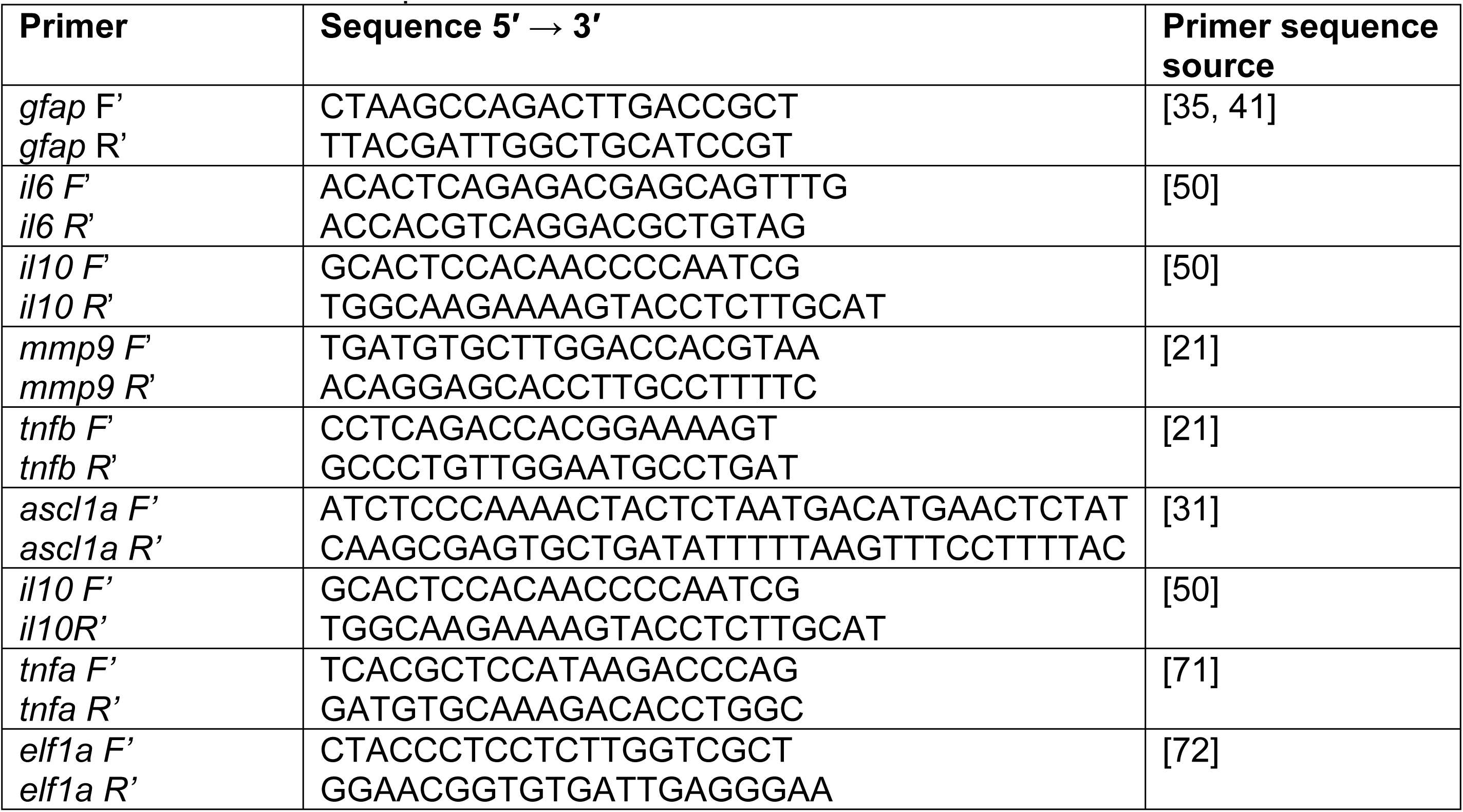
Primers used for qPCR.

### Graphing and Statistical Analysis

Graphing and statistics were performed in the R coding environment or using GraphPad Prism. For pairwise comparisons of quantitative data, either parametric (t-test for normally distributed data) or non-parametric Mann-Whitney U-tests, or for data with unequal variances Welch’s test, were performed. For proportion data, Fisher Exact test was used. For data with multiple factors, 2-way ANOVA, Kruskal-Wallis, or 3-way ANOVA test was used, followed by post-hoc pairwise comparisons when warranted, as indicated in the figure legends. A p-value <0.05 was considered statistically significant. Statistically significant differences are annotated in the figures and stated in the figure legends.

## Acknowledgements

We thank Dr. Mikiko Nagashima (University of Michigan) for the kind gift of the *gnat2*:nfsb-mCherry transgenic line made in Rachel Wong’s laboratory (University of Washington). We thank the University of Idaho Imaging and Data Acquisition Core (IDAC, managed by the Institute of Modeling Complex Interactions/IMCI) and IIDS for support of our research. We thank Mae Dawson (undergraduate research student) for technical assistance.

## Bibliography

[1] Nagashima M, Barthel LK, Raymond PA. A self-renewing division of zebrafish Müller glial cells generates neuronal progenitors that require N-cadherin to regenerate retinal neurons. Development 2013; 140: 4510–4521.

[2] Bernardos RL, Barthel LK, Meyers JR, et al. Late-Stage Neuronal Progenitors in the Retina Are Radial Müller Glia That Function as Retinal Stem Cells. J Neurosci 2007; 27: 7028–7040.

[3] Thummel R, Kassen SC, Enright JM, et al. Characterization of Müller glia and neuronal progenitors during adult zebrafish retinal regeneration. Exp Eye Res 2008; 87: 433–444.

[4] Fausett BV, Goldman D. A Role for α1 Tubulin-Expressing Müller Glia in Regeneration of the Injured Zebrafish Retina. J Neurosci 2006; 26: 6303–6313.

[5] Fimbel SM, Montgomery JE, Burket CT, et al. Regeneration of Inner Retinal Neurons after Intravitreal Injection of Ouabain in Zebrafish. J Neurosci 2007; 27: 1712–1724.

[6] Thummel R, Kassen SC, Montgomery JE, et al. Inhibition of Müller glial cell division blocks regeneration of the light-damaged zebrafish retina. Dev Neurobiol 2008; 68: 392–408.

[7] McGinn TE, Mitchell DM, Meighan PC, et al. Restoration of Dendritic Complexity, Functional Connectivity, and Diversity of Regenerated Retinal Bipolar Neurons in Adult Zebrafish. J Neurosci 2018; 38: 120–136.

[8] McGinn TE, Galicia CA, Leoni DC, et al. Rewiring the Regenerated Zebrafish Retina: Reemergence of Bipolar Neurons and Cone-Bipolar Circuitry Following an Inner Retinal Lesion. Frontiers Cell Dev Biology 2019; 7: 95.

[9] Sherpa T, Lankford T, McGinn TE, et al. Retinal regeneration is facilitated by the presence of surviving neurons. Dev Neurobiol 2014; 74: 851–876.

[10] Barrett LM, Mitchell DM, Meighan PC, et al. Dynamic functional and structural remodeling during retinal regeneration in zebrafish. Front Mol Neurosci 2022; 15: 1070509.

[11] Abraham E, Hartmann H, Yoshimatsu T, et al. Restoration of cone-circuit functionality in the regenerating adult zebrafish retina. Dev Cell 2024; 59: 2158–2170.e6.

[12] Hammer J, Röppenack P, Yousuf S, et al. Visual Function is Gradually Restored During Retina Regeneration in Adult Zebrafish. Front Cell Dev Biol 2022; 9: 831322.

[13] Mitchell DM, Lovel AG, Stenkamp DL. Dynamic changes in microglial and macrophage characteristics during degeneration and regeneration of the zebrafish retina. J Neuroinflamm 2018; 15: 163.

[14] White DT, Sengupta S, Saxena MT, et al. Immunomodulation-accelerated neuronal regeneration following selective rod photoreceptor cell ablation in the zebrafish retina. Proc National Acad Sci 2017; 114: E3719–E3728.

[15] Bludau O, Weber A, Bosak V, et al. Inflammation is a critical factor for successful regeneration of the adult zebrafish retina in response to diffuse light lesion. Front Cell Dev Biol 2024; 12: 1332347.

[16] Iribarne M, Hyde DR. Different inflammation responses modulate Müller glia proliferation in the acute or chronically damaged zebrafish retina. Front Cell Dev Biol 2022; 10: 892271.

[17] Fischer AJ, Zelinka C, Gallina D, et al. Reactive microglia and macrophage facilitate the formation of Müller glia-derived retinal progenitors. Glia 2014; 62: 1608–1628.

[18] Song P, Parsana D, Singh R, et al. Photoreceptor regeneration occurs normally in microglia-deficient irf8 mutant zebrafish following acute retinal damage. Sci Rep 2024; 14: 20146.

[19] Xu H, Cao L, Chen Y, et al. Single-cell RNA sequencing reveals the heterogeneity and interactions of immune cells and Müller glia during zebrafish retina regeneration. Neural Regen Res 2024; 20: 3635–3648.

[20] Conedera FM, Pousa AMQ, Mercader N, et al. Retinal microglia signaling affects Müller cell behavior in the zebrafish following laser injury induction. Glia 2019; 67: 1150–1166.

[21] Silva NJ, Nagashima M, Li J, et al. Inflammation and matrix metalloproteinase 9 (Mmp-9) regulate photoreceptor regeneration in adult zebrafish. Glia 2020; 68: 1445–1465.

[22] Fogerty J, Song P, Boyd P, et al. Notch Inhibition Promotes Regeneration and Immunosuppression Supports Cone Survival in a Zebrafish Model of Inherited Retinal Dystrophy. J Neurosci 2022; 42: 5144–5158.

[23] D’Orazi FD, Suzuki SC, Darling N, et al. Conditional and biased regeneration of cone photoreceptor types in the zebrafish retina. J Comp Neurol 2020; 528: 2816–2830.

[24] Thomas JL, Nelson CM, Luo X, et al. Characterization of multiple light damage paradigms reveals regional differences in photoreceptor loss. Exp Eye Res 2012; 97: 105–116.

[25] Zhang Z, Hou H, Yu S, et al. Inflammation-induced mammalian target of rapamycin signaling is essential for retina regeneration. Glia 2020; 68: 111–127.

[26] García-García D, Vidal-Gil L, Parain K, et al. Neuroinflammation as a cause of differential Müller cell regenerative responses to retinal injury. Sci Adv 2024; 10: eadp7916.

[27] Rumford JE, Grieshaber A, Lewiston S, et al. Forced MyD88 signaling in microglia impacts the production and survival of regenerated retinal neurons. Front Cell Dev Biol 2024; 12: 1495586.

[28] Todd L, Finkbeiner C, Wong CK, et al. Microglia Suppress Ascl1-Induced Retinal Regeneration in Mice. Cell Reports 2020; 33: 108507.

[29] Morales M, Findley AP, Mitchell DM. Intercellular contact and cargo transfer between Müller glia and to microglia precede apoptotic cell clearance in the developing retina. Development; 151. Epub ahead of print 2024. DOI: 10.1242/dev.202407.

[30] Harada T, Harada C, Kohsaka S, et al. Microglia-Müller glia cell interactions control neurotrophic factor production during light-induced retinal degeneration. J Neurosci : Off J Soc Neurosci 2002; 22: 9228–36.

[31] Mitra S, Devi S, Lee M-S, et al. Vegf signaling between Müller glia and vascular endothelial cells is regulated by immune cells and stimulates retina regeneration. Proc National Acad Sci 2022; 119: e2211690119.

[32] Raghavan D, Jeakle O, Berry Y, et al. Microglia response and function in a chronic model of photoreceptor damage. Front Cell Dev Biol 2025; 13: 1699271.

[33] Blume ZI, Lambert JM, Lovel AG, et al. Microglia in the developing retina couple phagocytosis with the progression of apoptosis via P2RY12 signaling. Dev Dynam 2020; 249: 723–740.

[34] Ellett F, Pase L, Hayman JW, et al. mpeg1 promoter transgenes direct macrophage-lineage expression in zebrafish. Blood 2011; 117: e49–e56.

[35] Mitchell DM, Sun C, Hunter SS, et al. Regeneration associated transcriptional signature of retinal microglia and macrophages. Sci Rep-uk 2019; 9: 4768.

[36] Leach LL, Hanovice NJ, George SM, et al. The immune response is a critical regulator of zebrafish retinal pigment epithelium regeneration. Proc Natl Acad Sci 2021; 118: e2017198118.

[37] Gallina D, Zelinka C, Fischer AJ. Glucocorticoid receptors in the retina, Müller glia and the formation of Müller glia-derived progenitors. Development 2014; 141: 3340–3351.

[38] Gallina D, Zelinka CP, Cebulla CM, et al. Activation of glucocorticoid receptors in Müller glia is protective to retinal neurons and suppresses microglial reactivity. Exp Neurol 2015; 273: 114–125.

[39] Emmerich K, White DT, Kambhampati SP, et al. Nanoparticle-based targeting of microglia improves the neural regeneration enhancing effects of immunosuppression in the zebrafish retina. Commun Biol 2023; 6: 534.

[40] Shiau CE, Kaufman Z, Meireles AM, et al. Differential Requirement for irf8 in Formation of Embryonic and Adult Macrophages in Zebrafish. Plos One 2015; 10: e0117513.

[41] Thiel WA, Blume ZI, Mitchell DM. Compensatory engulfment and Müller glia reactivity in the absence of microglia. Glia 2022; 70: 1402–1425.

[42] Oosterhof N, Kuil LE, Linde HC van der, et al. Colony-Stimulating Factor 1 Receptor (CSF1R) Regulates Microglia Density and Distribution, but Not Microglia Differentiation In Vivo. Cell Reports 2018; 24: 1203–1217.e6.

[43] Leach LL, Gonzalez RG, Jayawardena SU, et al. Macrophage/microglia-dependent mechanisms drive retinal pigment epithelium regeneration in zebrafish. Cell Rep 2025; 44: 116325.

[44] D’Orazi FD, Zhao X-F, Wong RO, et al. Mismatch of Synaptic Patterns between Neurons Produced in Regeneration and during Development of the Vertebrate Retina. Curr Biol 2016; 26: 2268–2279.

[45] Rovira M, Miserocchi M, Montanari A, et al. Zebrafish Galectin 3 binding protein is the target antigen of the microglial 4C4 monoclonal antibody. Dev Dyn 2023; 252: 400–414.

[46] Bailey TJ, Fossum SL, Fimbel SM, et al. The inhibitor of phagocytosis, O-phospho-l-serine, suppresses Müller glia proliferation and cone cell regeneration in the light-damaged zebrafish retina. Exp Eye Res 2010; 91: 601–612.

[47] Guyader DL, Redd MJ, Colucci-Guyon E, et al. Origins and unconventional behavior of neutrophils in developing zebrafish. Blood 2008; 111: 132–141.

[48] Anderson SR, Roberts JM, Ghena N, et al. Neuronal apoptosis drives remodeling states of microglia and shifts in survival pathway dependence. Elife; 11. Epub ahead of print 2022. DOI: 10.7554/elife.76564.

[49] Elmore MRP, Najafi AR, Koike MA, et al. Colony-Stimulating Factor 1 Receptor Signaling Is Necessary for Microglia Viability, Unmasking a Microglia Progenitor Cell in the Adult Brain. Neuron 2014; 82: 380–397.

[50] Lu C, Hyde DR. Cytokines IL-1β and IL-10 are required for Müller glia proliferation following light damage in the adult zebrafish retina. Front Cell Dev Biol 2024; 12: 1406330.

[51] Nelson CM, Ackerman KM, O’Hayer P, et al. Tumor Necrosis Factor-Alpha Is Produced by Dying Retinal Neurons and Is Required for Müller Glia Proliferation during Zebrafish Retinal Regeneration. J Neurosci 2013; 33: 6524–6539.

[52] Bringmann A, Iandiev I, Pannicke T, et al. Cellular signaling and factors involved in Müller cell gliosis: Neuroprotective and detrimental effects. Prog Retin Eye Res 2009; 28: 423–451.

[53] Thomas JL, Ranski AH, Morgan GW, et al. Reactive gliosis in the adult zebrafish retina. Exp Eye Res 2016; 143: 98–109.

[54] Clark DT. Visual Responses in Developing Zebrafish (Brachydanio Rerio). University of Oregon, https://books.google.com/books?id=D_wHOAAACAAJ.

[55] Bilotta J, Saszik S, Sutherland SE. Rod contributions to the electroretinogram of the dark-adapted developing zebrafish. Dev Dyn 2001; 222: 564–570.

[56] Branchek T, Bremiller R. The development of photoreceptors in the zebrafish, Brachydanio rerio. I. Structure. J Comp Neurol 1984; 224: 107–115.

[57] Haug MF, Berger M, Gesemann M, et al. Differential expression of PKCα and -β in the zebrafish retina. Histochem Cell Biol 2019; 151: 521–530.

[58] Suzuki S, Kaneko A. Identification of bipolar cell subtypes by protein kinase C-like immunoreactivity in the goldfish retina. Vis Neurosci 1990; 5: 223–230.

[59] D’Orazi FD, Suzuki SC, Wong RO. Neuronal remodeling in retinal circuit assembly, disassembly, and reassembly. Trends Neurosci 2014; 37: 594–603.

[60] Saade CJ, Alvarez-Delfin K, Fadool JM. Rod Photoreceptors Protect from Cone Degeneration-Induced Retinal Remodeling and Restore Visual Responses in Zebrafish. J Neurosci 2013; 33: 1804–1814.

[61] Powell C, Cornblath E, Elsaeidi F, et al. Zebrafish Müller glia-derived progenitors are multipotent, exhibit proliferative biases and regenerate excess neurons. Sci Rep 2016; 6: 24851.

[62] Kei JNC, Currie PD, Jusuf PR. Fate bias during neural regeneration adjusts dynamically without recapitulating developmental fate progression. Neural Dev 2017; 12: 12.

[63] Lahne M, Brecker M, Jones SE, et al. The Regenerating Adult Zebrafish Retina Recapitulates Developmental Fate Specification Programs. Front Cell Dev Biol 2021; 8: 617923.

[64] Hanovice NJ, Leach LL, Slater K, et al. Regeneration of the zebrafish retinal pigment epithelium after widespread genetic ablation. Plos Genet 2019; 15: e1007939.

[65] George SM, Lu F, Rao M, et al. The retinal pigment epithelium: Development, injury responses, and regenerative potential in mammalian and non-mammalian systems. Prog Retin Eye Res 2021; 85: 100969.

[66] Konishi H, Okamoto T, Hara Y, et al. Astrocytic phagocytosis is a compensatory mechanism for microglial dysfunction. EMBO J 2020; 39: EMBJ2020104464.

[67] Beachum AN, Salazar G, Nachbar A, et al. Glia Preserve Their Own Functions While Compensating for Neighboring Glial Cell Dysfunction. Glia. Epub ahead of print 2025. DOI: 10.1002/glia.70072.

[68] Westerfield M. The zebrafish book. A guide for the laboratory use of zebrafish (Danio rerio). 5th Ed. University of Oregon Press, 2007.

[69] Meeker ND, Hutchinson SA, Ho L, et al. Method for isolation of PCR-ready genomic DNA from zebrafish tissues. BioTechniques 2007; 43: 610–614.

[70] Lovel AG, Mitchell DM. Axon Regeneration, Methods and Protocols. Methods Mol Biol 2023; 2636: 389–400.

[71] Tsarouchas TM, Wehner D, Cavone L, et al. Dynamic control of proinflammatory cytokines Il-1β and Tnf-α by macrophages in zebrafish spinal cord regeneration. Nat Commun 2018; 9: 4670.

[72] Kirchberger S, Shoeb MR, Lazic D, et al. Comparative transcriptomics coupled to developmental grading via transgenic zebrafish reporter strains identifies conserved features in neutrophil maturation. Nat Commun 2024; 15: 1792.

